# The RNA-DNA world and the emergence of DNA-encoded heritable traits

**DOI:** 10.1101/2023.03.14.532532

**Authors:** Suvam Roy, Supratim Sengupta

## Abstract

The RNA world hypothesis confers a central role to RNA molecules in information encoding and catalysis. Even though evidence in support of this hypothesis has accumulated from both experiments and computational modelling, the transition from an RNA world to a world where heritable genetic information is encoded in DNA remains an open question. Recent experiments show that both RNA and DNA templates can extend complementary primers using free RNA/DNA nucleotides, either non-enzymatically or in the presence of a replicase ribozyme. Guided by these experiments, we analyse protocellular evolution with an expanded set of reaction pathways made possible through the presence of DNA nucleotides. By encapsulating these reactions inside three different types of protocellular compartments, each subject to distinct modes of selection, we show how protocells containing DNA-encoded replicases in low copy numbers and replicases in high copy numbers can dominate the population. This is facilitated by a reaction that leads to auto-catalytic synthesis of replicase ribozymes from DNA templates encoding the replicase, after the chance emergence of a replicase through non-enzymatic reactions. Our work unveils a pathway for the transition from an RNA world to a mixed RNA-DNA world characterized by Darwinian evolution, where DNA sequences encode heritable phenotypes.

## 1 Introduction

The RNA world hypothesis states that a living system composed entirely of RNA may have emerged prior to the origin of the current DNA-protein world. The ability of RNA to store information [1, 2] and catalyse reactions [3, 4, 5] suggest that they may indeed have played a key role in the emergence of a self-sustaining chemical system capable of undergoing Darwinian evolution. However, the plausibility of such an RNA world relies on explaining how self-replicating protocells encapsulating RNA molecules first emerged from a primordial soup containing basic chemical building blocks. Several experiments have shown that both ribonucleotides [6, 7, 8] and deoxy-ribonucleotides [9, 10] can be spontaneously created in prebiotic environments. Non-enzymatic processes of concatenation and template-directed primer extension, albeit with [11, 12] or without [13] activated RNA nucleotides, along with environmental cycling [14, 15] give rise to long RNA polymers that can fold into complex secondary structures [16]. It seems plausible that the first ribozymes emerged by chance through such processes, as a result of extensive sampling of the sequence space through error-prone replication [17] in a prebiotic environment. Recent work has provided evidence of phenotype bias, indicating that certain secondary structures which appear most frequently while sampling sequences of fixed length happen to be the ones that are found in nature [18]. Remarkably, those structures can be generated by sampling a relatively small region of the sequence space, implying that the chance of emergence of a functional biomolecule may not be too small. However, the random creation of a few functional bio-molecules is not sufficient to guarantee their proliferation especially in a spatially open environment dominated by selfish parasitics equences. It is therefore imperative for template-directed replication processes to be encapsulated in compartments since such segregation can help in preventing parasites from overwhelming the system [19]. If distinct ribozymes are created with small probabilities inside a protocell during replication by the rolling circle mechanism [20, 21, 22, 23, 24], they can act to not only synergistically enhance each others’ formation but also increase the likelihood of creation of other functionally distinct ribozymes. This provides a pathway towards increasing functional complexity of the protocell and can allow for the proliferation of protocells containing a functionally diverse set of ribozymes [25]. However, such an evolution is not Darwinian in nature and the ribozymes generated cannot be considered as heritable traits since they appear by chance during the error-prone replication process and are also prone to degradation. How then is it possible to create a self-sustained chemical system that does not rely on random ribozyme production for its survival? A possible resolution to this problem in the RNA world was proposed by Takeuchi *et al*. [26] who showed that genome-like strands can appear inside vesicles due to symmetry-breaking between two complementary strands of a self-replicating ribozyme. One strand would then act as a genome and its complementary strand as an enzyme. Since there is very little free-energy difference between a sequence and its complement, it seems unlikely that the reverse complement of a template strand would fold into a complex secondary structure, that is characteristic of a ribozyme. We believe the answer to the riddle of creating a self-sustained chemical replicator lies in the appearance of information-encoding DNA templates that act like primitive pseudo-genes. A collection of such pseudo-genes, each encoding a different functional phenotype, would then constitute a primitive albeit fragmented genome. However, unlike ribonucleotides [27, 28, 29], spontaneous polymerization of free DNA nucleotides into long DNA strands have not yet been demonstrated by experiments. Therefore the transition to a DNA world must have been facilitated by RNA. Indeed, experiments on mixed RNA-DNA template-primer systems have shown that RNA is capable of extending DNA primers both non-enzymatically [30] and under the action of polymerase ribozymes [31, 32, 33]. Non-enzymatic and enzymatic reverse transcription of RNA strands might have been the primary mechanism of DNA strand creation in a primordial world. These results [31, 32, 33] are particularly significant because they suggest that the transition from an RNA world to a DNA-protein world passed through an intermediate epoch where life may have been based on DNA genomes and ribozymes that took on the role of proteins.

In this work, we show how the presence of DNA nucleotides increases the space of possible reactions and opens up the possibility of creating single-stranded DNA sequences that can act as information-encoding templates. Even though the error-prone non-enzymatic template-directed replication process produces mostly useless sequences (parasites), we assume that biomolecules like replicase ribozymes can be created with small probabilities. The replication of RNA templates using DNA nucleotides can also create, with a small probability, the DNA sequences encoding the replicase ribozymes. Those DNA sequences can act as templates, whose accurate, auto-catalytic replication with RNA nucleotides can create more copies of the replicase. Such templates along with the replicase can speed up the replication process inside a protocell which in-turn ensures rapid growth in the number of strands and enhanced likelihood of creation of new replicases and templates. A population of protocells can eventually evolve to increase the number of protocells containing both replicase-encoding templates and replicases thereby marking the onset of Darwinian evolution characterized by the encoding of a heritable trait like the replicase ribozyme in DNA sequences. The nature of the protocellular compartment determines how likely it is for such protocells to dominate the population under reasonable conditions. Proliferation of such protocells is observed for coacervate droplets, water-in-oil droplets as well as vesicular compartments. However, coacervate droplets and water-in-oil droplets, allow for the dominance of such protocells under diverse conditions that are even robust to decrease in the error threshold for accurate replication. Intriguingly, in our model, evolution leads to the spontaneous emergence of proto-cells that contain the replicase-encoding DNA templates in low copy numbers and replicases in high copy numbers. Our work shows how the creation of DNA-encoded heritable phenotypes through RNA-templated replication can lead to the emergence of protocells capable of undergoing Darwinian evolution.

## 2 Methods

The reactions inside each protocell depends on the nature of the components present in it. The presence of DNA nucleotides allows for the non-enzymatic replication of RNA templates using deoxy-ribonucleotides [30, 34] leading to the creation of DNA strands. The number of strands inside a protocell initially grow through non-enzymatic template-directed replication based on the reactions described below (see section 2 of the electronic supplementary material for details of estimation of the replication rates). Enzymatic reactions are activated once a replicase emerges as a result of a chance non-enzymatic replication event. Table-1 gives both the non-enzymatic and enzymatic replication rates for the various reactions and Supplementary Fig-S1 gives a pictorial representation of the different processes considered. The evolution of a population of such protocells occurs due to proto-cellular growth followed by division and selection in a manner that depends on the nature of the encapsulating compartment.

**Table 1:** Non-enzymatic and enzymatic replication rates for a template of length L=200 nt inside a protocell of radius 57 nm along with the corresponding percentage errors (*ϵ* = 100 *× er/L*). *r* = *RNA, d* = *DNA, T* = *template, R* = *replicase*. 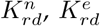 represent non-enzymatic and enzymatic replication rates from RNA to DNA etc.

| Reaction | Rate ($h^{-1}$) | Error (%) | Reference |
| --- | --- | --- | --- |
| $K_{rr}^n$ | $0.00255 T$ | $\sim 36\%$ | [17, 35] |
| $K_{rd}^n$ | $0.001646 T$ | High | [30] |
| $K_{dr}^n$ | $0.000823 T$ | High | [30] |
| $K_{dd}^n$ | $0.00255 T$ | $\sim 32\%$ | [17, 34] |
| $K_{rr}^e$ | $0.46447 R$ | $\sim 8\%$ | [31, 32] |
| $K_{rd}^e$ | $0.226 R$ | $\sim 3\%$ | [32] |
| $K_{dr}^e$ | $0.1434 R$ | $\sim 3\%$ | [33] |
| $K_{dd}^e$ | 0 | NA | [32] |

### 2.1 Reactions

Initially, we consider protocells containing RNA templates *T*_*r*_ only. Non-enzymatic replication of *T*_*r*_ by RNA nucleotides (with a rate 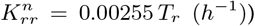 will mostly create RNA parasites (P) (which are non-functional sequences that are neither templates nor ribozymes), due to the high error-rates of non-enzymatic reactions. However, following the argument given in the Introduction, we assume that replicase ribozymes (R) can be created with a small probability (*p*_*R*_) during the error-prone, non-enzymatic replication process (see reaction (1)). The chance creation of a replicase eventually leads to the emergence of a more complex reaction network as indicated by the blue arrows in Fig-1. We also assume unlimited monomer supply by the environment to keep the rates independent of monomer concentrations. *P/R* on the RHS of reactions 1 indicates that both parasites (*P*) and replicases (*R*) can be produced albeit with different probabilities that are indicated in brackets.

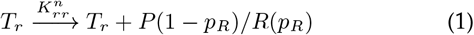

**Figure 1.**
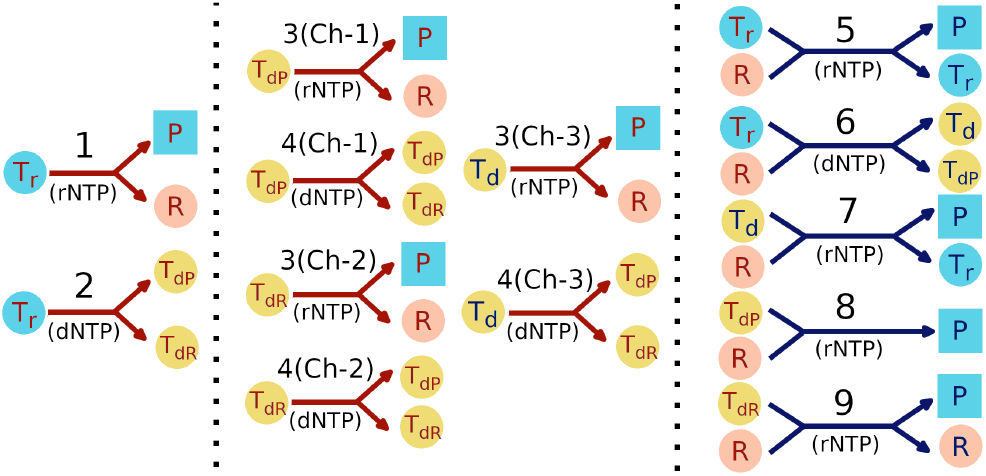
Pictorial representation of the reaction network inside a protocell. Red (blue) arrows indicate non-enzymatic (enzymatic) reactions. Parasites are indicated by square filled boxes whereas all other reactants and products are indicated by filled circles. Light blue filled circles denote RNA templates, peach colour filled circles denote replicases and yellow filled circles denote DNA templates. The type of monomer used in each reaction is indicated in brackets with rNTP and dNTP indicating RNA and DNA nucleotides respectively. The left panel shows the reactions possible initially in presence of RNA templates only. The middle panel shows the non-enzymatic reactions possible after the emergence of DNA templates. Ch-1, Ch-2 and Ch-3 denote 3 different channels for non-enzymatic replication of the 3 different types of DNA templates. The right panel (blue arrows) shows the enzymatic reactions possible upon emergence of a replicase ribozyme. The numbers 1-9 correspond to the reaction numbers specified in the main text. Arrows with multiple arrowheads denote the possible products of a reaction.

For the RNA world to be viable, sustained creation of replicases that can catalyse replication of template strands is essential. Non-enzymatic or enzymatic replication of a replicase, whose catalytic ability arises from its complex folded structure, is extremely difficult to achieve because folded segments can block replication of such RNA sequences. So far, the only way this can be achieved is by using trimer building blocks instead of monomers [36]. However, that process requires an abundance of trimers that are an exact complement of triplet nucleotides that make up the structured template. The abundance of such trimers, created via spontaneous concatenation of free monomers in a primordial RNA world, is likely to be much smaller than the abundance of monomers. Therefore the feasibility of replicating ribozyme sequences from structured templates in prebiotic scenarios remains questionable. This problem can be avoided if ribozymes are encoded in DNA sequences. We calculated the folding free energies of different RNA sequences and their DNA counterparts, and observed that free energies of the DNA strands are ∼ 3.4 times larger than the free energies of their RNA counterparts [37, 38, 39]. Therefore DNA strands are less likely to fold into complex secondary structures and more likely to act as templates.

The enzymatic replication of such ribozyme-encoding DNA templates using RNA nucleotides [33] can then provide a higher fidelity pathway for creation of those ribozymes. Recent experiments showing RNA templated DNA synthesis [32] and DNA templated RNA synthesis [33] seems to suggest the plausibility of such a scenario of sustained ribozyme creation.

Non-enzymatic replication of *T*_*r*_ by DNA nucleotides will create DNA templates, the majority of which will be non-functional since they will encode RNA parasites (*T*_*dP*_). However, such replication processes can also create, with a small probability *p*_*R*_, DNA templates (*T*_*dR*_) that encode replicase ribozymes. The probabilities *p*_*R*_ and (1 − *p*_*R*_) for creation of *T*_*dR*_ and *T*_*dP*_ respectively are indicated in brackets in the reaction below.

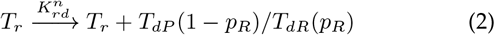

Non-enzymatic replication of DNA templates *T*_*d*_ (which is the DNA analog of *T*_*r*_), *T*_*dP*_ and *T*_*dR*_ by RNA nucleotides will create parasites and replicases 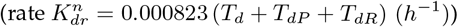. Similarly non-enzymatic replication of DNA templates *T*_*d*_, *T*_*dP*_ and *T*_*dR*_ by DNA nucleotides can create *T*_*dP*_ and *T*_*dR*_ (rate 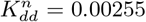 (*T*_*d*_ + *T*_*dP*_ + *T*_*dR*_)(*h*^−1^)) . *T*_*d*_*/T*_*dP*_ */T*_*dR*_ on the LHS of reactions 3-9 indicate that any one of them is used as a template and the corresponding template is therefore also present on the RHS of the reactions.

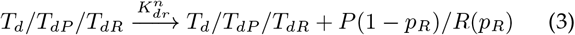

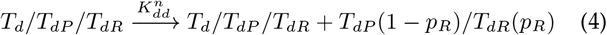

We assume the probability of *R* and *T*_*dR*_ creation to be same for all DNA templates *T*_*d*_, *T*_*dR*_ and *T*_*dP*_, because non-enzymatic replication processes have very high nucleotide misincorporation rates (see Table-1). Therefore, such replication processes effectively leads to the creation of randomly sampled 200-mer sequences that are likely to be uncorrelated with the sequence of the underlying templates. It then seems reasonable to expect that the chances of non-enzymatic creation of *R* and *T*_*dR*_ would be independent of the nature of the template.

Initially, the processes inside a protocell are driven only by non-enzymatic reactions. However, upon the chance emergence of replicase ribozymes, enzymatic replication of *T*_*r*_ [31] by RNA nucleotides (see reaction 5) can create new copies of the template as the accuracy of the enzymatic reactions is much higher compared to the non-enzymatic reactions. Nevertheless, parasites will continue to be created, since even enzymatic replication in a primordial world is error-prone (Table-1) in the absence of proof-reading mechanisms. We define a function *f* (*ϵ*) = 1*/*(1 + *exp*(*ϵ* − *e*_*T*_)), where *e*_*T*_ is the error-threshold, to quantify the likelihood of accurate replication under the action of a replicase ribozyme. This depends on the percentage error (*ϵ*) during such enzymatic replications (see Table-1). Fig. S2 shows the variation of this probability of accurate replication for the three types of enzymatic replication processes with different values of the error threshold *e*_*T*_. Enzymatic replication of *T*_*r*_ by DNA nucleotides [32] will similarly create (see reaction 6) a DNA version *T*_*d*_ of the template *T*_*r*_ or DNA template *T*_*dP*_ encoding a parasitic sequence, with different likelihoods modulated by the function *f* (*ϵ*_*rd*_).

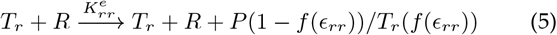

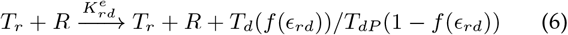

The three types of DNA templates in our model allow for three different enzymatic replications of DNA templates (with a rate 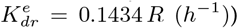. Enzymatic replication of *T*_*d*_ by RNA nucleotides will create primarily parasites but can also produce an RNA template *T*_*r*_ with probability *f* (*ϵ*_*dr*_) (see reaction 7). In the case of *T*_*dP*_, enzymatic replication by RNA nucleotides will create only parasites (see reaction 8). Finally, auto-catalytic replication of *T*_*dR*_ by RNA nucleotides can recreate a replicase depending on the error-threshold.

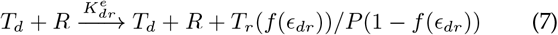

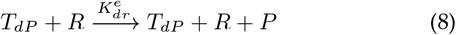

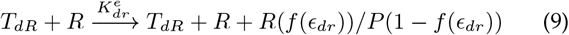

Reaction 9 marks the onset of one way information flow from the DNA template sequence to the replicase it encodes and can be considered to be a manifestation of a primitive central dogma. Fig-1 provides a pictorial representation of the nine types of replication reactions that can occur inside a protocell. Additionally, we also consider degradation of both RNA and DNA molecules. As DNA is more stable than RNA, the degradation rate of DNA (*h*_*d*_) is taken to be lower than RNA (*h*_*r*_). We use *h*_*r*_ = 0.0008 *h*^−1^, *h*_*d*_ = 0.00008 *h*^−1^ for all simulations.

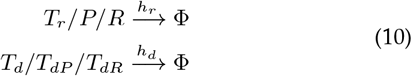

The differential equations determining the time evolution of the abundances of the six different types of molecules, as a result of reactions (1)-(10), are given in the *electronic supplementary material*.

### 2.2 Protocellular compartments and population evolution

We considered N=400 protocells each initially containing 10 RNA templates *T*_*r*_. To account for variation between different proto-cells, we assumed that the templates inside different protocells have different replicase creation probabilities, with the probability taken randomly from the range (0.5 *p*_*R*_, 1.5 *p*_*R*_). The two key parameters in our model are the creation probability (*p*_*R*_) of *R/T*_*dR*_ via error-prone, non-enzymatic replication and the error-threshold (*e*_*T*_) in case of enzymatic reactions. We varied these 2 parameters and carried out stochastic simulations of the evolving population. The dynamics of the evolving population can be best understood by tracking the average fraction of replicase per protocell, fraction of protocells containing both replicase *R*, the corresponding DNA strand (*T*_*dR*_) encoding it and the relative propensity of reaction 9 (see *electronic supplementary material* for details on how these quantities were estimated.)

The three different types of protocellular compartments in our model are distinguished by the distinct selection mechanisms they undergo during the evolution of the protocellular population. Fig-2 gives a schematic representation of the different types of selection mechanisms.

**Figure 2.**
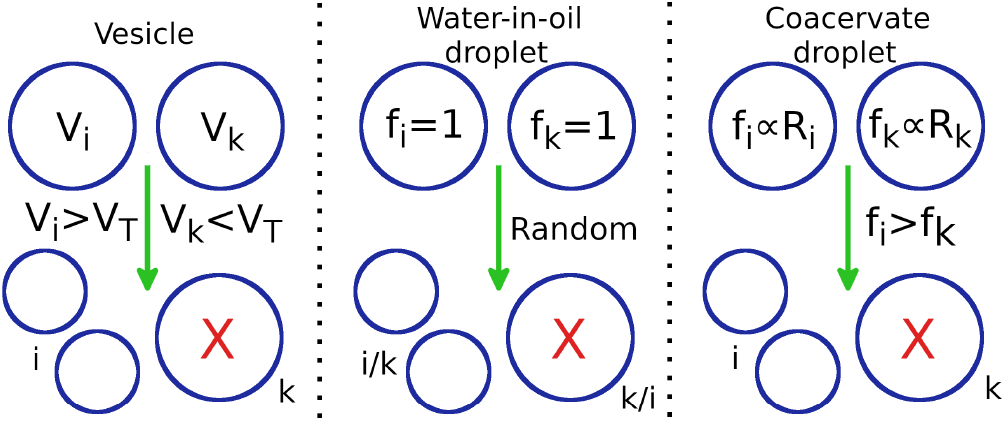
Schematic representation of three different modes of protocellular competition: *i, k* corresponds to the *i*^*th*^ and *k*^*th*^ protocell. *V*_*n*_ denotes the total number of strands inside the *n*^*th*^ protocell, *V*_*T*_ is the upper limit of *V* and *f* is the fitness of the protocell. *Vesicles:* if *V*_*i*_ exceeds the upper limit *V*_*T*_, while *V*_*k*_ *< V*_*T*_, *i* will divide into 2 daughter vesicles and *k* is eliminated. *Water-in-oil droplets: i* and *k* are equally likely to eliminate each other through a random selection process. The surviving droplet divides into two daughter droplets. *Coacervate droplets:* if *i* contains larger number (or fraction) of ribozymes (*R*_*i*_) compared to *k*; *f*_*i*_ *> f*_*k*_ and *i* is more likely to eliminate *k* and divide into two daughter protocells.

In the first scenario, our choice of protocellular compartments made out of water-in-oil droplets was inspired by recent RNA host-parasite experiments [40, 41, 42, 43, 44]. In the experiments, the protocellular compartments are subjected to periodic washout-mixing cycles where a fraction of droplets are randomly removed from the population followed by supply of empty compartments which are then mixed with existing compartments leading to random redistribution of components from the filled to empty compartments. We model this system by considering pairwise competition between protocells that is initiated whenever the number of strands inside all droplets lie in the range (*V /*5, *V*). During such competition, each droplet competes with another droplet chosen at random from the population; one of those two is then randomly selected to be eliminated while the other divides into 2 daughter protocells with components of the surviving protocell being randomly distributed between the two daughter droplets.

As another alternative, we consider the competition between protocells whose fitness is determined by functional molecules ike ribozymes encapsulated in it. Such genotype-phenotype coupling is an essential feature of Darwinian evolution since internal functional components can provide a fitness advantage thereby ensuring preferential selection of such protocells. Coacervate droplets formed by liquid-liquid phase separation [45, 46, 47, 48] can be considered to be an ideal candidate for those types of protocells. Such membraneless droplets can selectively partition biomolecules, sequester long RNA molecules [49], undergo growth and division [50] under environmental cycling [51], support key prebiotic processes like catalysis [45], template-directed primer extension [52] and ligation [53] and show enhanced catalytic activity of encapsulated ribozymes [45, 53, 54]. Moreover, the presence of active ribozymes inside coacervate droplets have been shown to modulate droplet properties [53, 55] thereby establishing a genotype-phenotype linkage and conferring on such droplets a potential fitness advantage under certain environmental conditions.

Therefore in our coacervate droplet model we consider a competition similar to the water-in-oil droplet model but with selection dependent not just on the number of strands but on the fraction of replicase ribozymes inside the droplet. This amounts to defining a fitness function *f*_*i*_ for each droplet involved in pairwise competition with 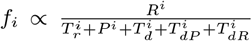. The droplet selected on the basis of its fitness divides into two daughter droplets, one of which replaces the droplet eliminated during pairwise competition.

Finally, vesicular compartments with lipid bilayer membranes were also considered. The volume of the vesicle is determined by the number but not the nature of strands inside it and vesicle division occurs when the number of strands reach an upper bound set to *V* = 1000. During the division process, the components of the vesicle are randomly distributed among the two daughter vesicles while an existing vesicle is eliminated with a probability proportional to the difference in the number of strands between itself and the dividing vesicle. This results in the preferential elimination of smaller vesicles [56] while keeping the population size fixed.

These three different types of competition between protocells leads to distinct conditions for the dominance of protocells containing a replicase ribozyme whose synthesis is brought about primarily through template-directed, replicase-catalysed replication of a DNA sequence encoding the enzyme.

## 3 Results

The presence of DNA nucleotides inside a single protocell can enhance the process of replicase creation and ensure sustained growth of strands. When enzymatic reactions are activated due to the chance emergence of a replicase, the presence of even a single DNA template ensures rapid growth of both DNA and RNA strands that further facilitate replicase production (see section 5 of electronic supplementary material for details). In the sub-sections below, we describe how the encapsulation of strands in three different types of protocellular compartments affect the outcome of competition between protocells in an evolving population.

### 3.1 Water-in-Oil droplet model

Fig-3 shows the heatmaps at equilibrium for the average fraction of replicase per droplet (panel A), fraction of droplets containing both replicase *R &* the DNA template (*T*_*dR*_) encoding it (panel B); and the relative propensity of reaction 9 (panel C) which provides a measure of the efficacy of replicase creation from its encoded DNA template. A very low non-enzymatic probability of replicase creation (*p*_*R*_ *<* 0.002) is not favourable for ensuring the presence of both *R* and *T*_*dR*_ in majority of droplets (see lower part of Fig-3(B)). A low value of *p*_*R*_ leads to low copy numbers of both *R* and *T*_*dR*_ through non-enzymatic processes which primarily lead to the creation of parasites (*P*) or parasite encoding DNA (*T*_*dP*_). The enzymatic creation of replicases through reaction 9 is suppressed since that pathway depends on the easy availability of both *R* and *T*_*dR*_. Even when an increase in the error threshold increases the probability of creation of *T*_*r*_ and *T*_*d*_ through reactions 5-7, it does not have an impact because of the suppression in creation of *R* and *T*_*dR*_ ensures that the reactant concentrations for 5-7 remain quite low. Even if the probability of non-enzymatic creation of replicases is relatively high (upper left region of Fig-3), low enzymatic replication fidelity due to very low error thresholds make parasite creation through enzymatic reaction channels 5-9 more likely, leading to lower average replicase fraction per droplet (see upper left region of Fig-3(A)).

**Figure 3.**
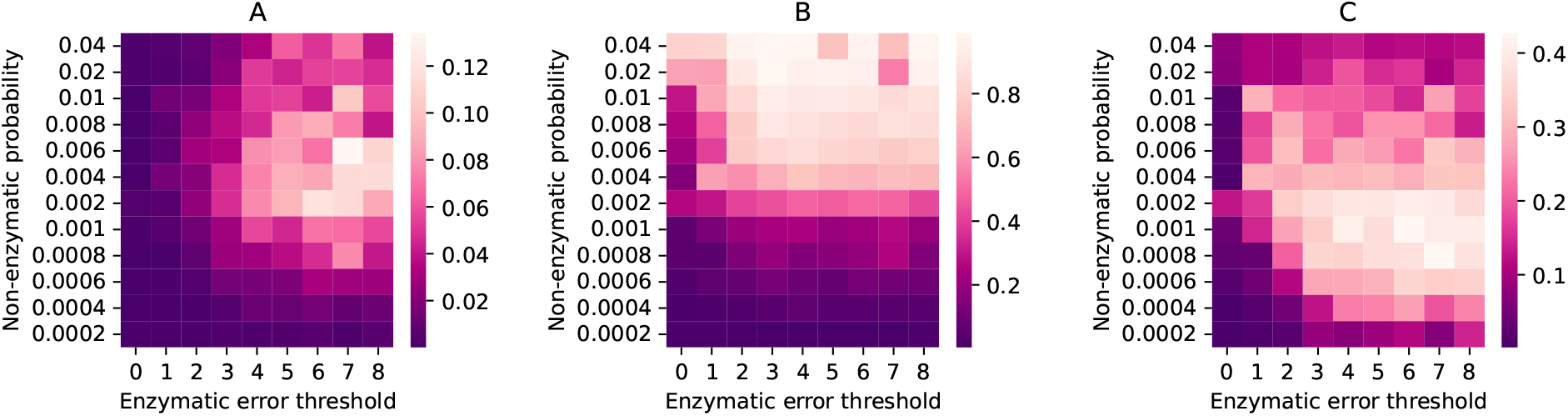
Stochastic simulation of a population of water-in-oil droplets containing strands. Heatmaps for **A:** average fraction of replicase ribozymes per droplet; **B:** fraction of droplets containing both replicase (*R*) and replicase encoding DNA template (*T*_*dR*_); **C:** average reaction propensity of reaction 9 per droplet; with different non-enzymatic ribozyme creation probabilities and error thresholds of enzymatic replications. The heatmaps are generated by taking both ensemble average and time average of the quantities at equilibrium.

When the non-enzymatic replicase creation probability (*p*_*R*_) as well as the error threshold (*e*_*T*_) are high, replicases are produced with a high likelihood both non-enzymatically and enzymatically since the replicase phenotype is more robust to replication errors. However, the resultant replicases are predominantly used to catalyze reactions 5-7 as evident from the higher reaction propensities of these reactions in the upper right region of Fig. S5(B). Replicases available for catalyzing their own formation are relatively fewer and reaction 9 is sub-dominant (see upper right region of Fig-3(C)). Moreover, since *T*_*r*_ and *T*_*d*_ are created with higher fidelity in this region of parameter space, the relative fraction of R is comparatively low as seen in Fig-3(A). In contrast, reaction 9 is dominant for moderately low non-enzymatic replicase creation probability and moderate to high error-threshold (see centre-right region of Fig-3(C)) leading to a comparatively larger average fraction of replicases per protocell. Fig-4 shows the time evolution of the average number of different strands (Fig-4(A)) for a point lying in this region. The percentage of droplets containing both *R* and *T*_*dR*_ (DNA template encoding *R*) is ∼ 48% (Fig-4(B)) and the average propensity of reaction 9, responsible for replicase creation using a DNA template encoding the replicase, is large (Fig-4(C)). Even though parasites continue to be formed, primarily through reaction 8, efficient creation of replicases both non-enzymatically and enzymatically is sufficient to ensure that close to a majority of protocells contain both *R* and *T*_*dR*_. The continuous regeneration of both *R* and *T*_*dR*_ can be sustained in the population even if the non-enzymatic reactions are eventually switched off after the onset of enzymatic replication of *R* (see Fig. S6). Intriguingly, we also find symmetry breaking between the number of *T*_*dR*_ and *R* with the average number of former and latter being ∼ 1 per droplet and ∼ 100 per droplet respectively (see Fig-4(A)). The crucial role played by reaction 9 in sustaining replicase formation and aiding the proliferation of droplets containing both *R* and *T*_*dR*_ is evident from Fig. S7 which shows the outcome of switching off reaction 9. Doing so prevents the asymmetry in copy numbers from emerging, drastically reduces the average number of both *R* and *T*_*dR*_ to below one per droplet and the percentage of droplets containing both *R* and *T*_*dR*_ to *<* 20 % of the population. This indicates that replicase creation through non-enzymatic processes alone cannot drive the evolution of the protocellular population towards increasing functional complexity.

**Figure 4.**
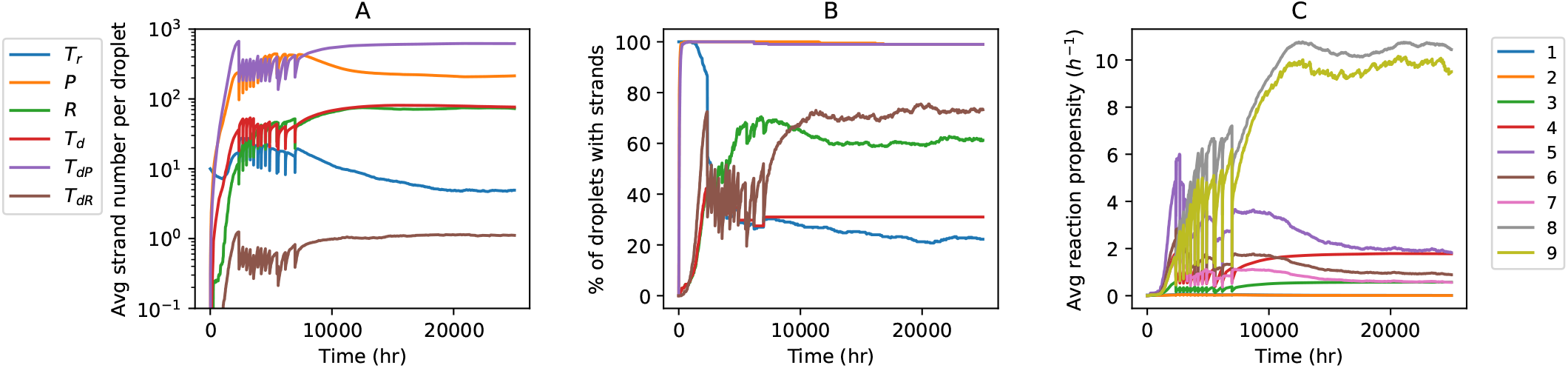
Stochastic simulation of a population of water-in-oil droplets (*p*_*R*_ = 0.002, *e*_*T*_ = 4). Time evolution of the **A:** Average number of different types of strands per droplet; **B:** Percentage of droplets containing different types of strands; **C:** Average reaction propensities of the 9 types of reactions per droplet. Panels **A** and **B** have the same legends.

### 3.2 Coacervate droplet model

In the case of coacervate droplets, selection pressure acts in favour of droplets containing more ribozymes. The region of parameter space where the fraction of droplets containing both *R & T*_*dR*_ exceeds ∼ 50% also overlaps with the region where the reaction propensity of 9 is large (Fig-5) and is characterized by moderately low values (≥𝒪 (10^−3^)) of non-enzymatic creation probability of a replicase. *T*_*dR*_ creation inside coacervate droplets can be amplified by harnessing the products (*T*_*d*_ and *T*_*dP*_) of the replicase catalysed reaction 6 and utilizing them to drive the non-enzymatic reactions 3-4 occurring through reaction channels Ch-1 and Ch-3. Hence, replicase creation through reaction 9 leads to a positive feedback that enhances the propensity of non-enzymatic reactions and induces the creation of even more replicases. Droplets which exploit such positive feedback loops resulting from creation of *R & T*_*dR*_ will be preferentially selected (see Fig. S8, Fig. S9(B) and the text below it in the electronic supplementary material) over droplets containing only parasites and can flourish even after non-enzymatic reactions are turned off (Fig. S10). The symmetry breaking between *R & T*_*dR*_ is also observed in this region (Fig. S9(A)), with the asymmetry between DNA templates and replicases vanishing in the absence of reaction 9 (see Fig. S11).

**Figure 5.**
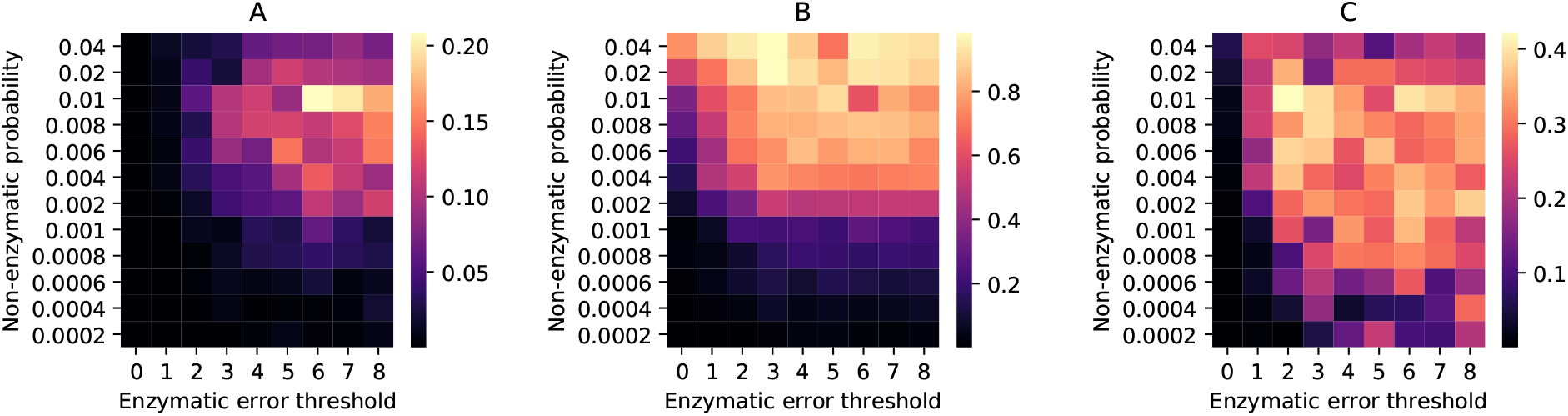
Stochastic simulation of a population of coacervate droplets containing strands. Heatmaps for **A:** average fraction of replicase ribozymes per droplet; **B:** fraction of droplets containing both replicase (*R*) and replicase encoding DNA template (*T*_*dR*_); **C:** average reaction propensity of reaction 9 per droplet; with different non-enzymatic ribozyme creation probabilities and error thresholds of enzymatic replications. The heatmaps are generated by taking both ensemble average and time average of the quantities at equilibrium.

### 3.3 Vesicle model

For such compartments, selection of a protocell depends only on the number but not on the type of strands in it, unlike the previous case. Since replicase formation is favoured both non-enzymatically (through reactions 1 and 3) and enzymatically (through reaction 9) for high *p*_*R*_ and high *e*_*T*_ respectively, the fraction of replicase per protocell increases with the non-enzymatic replicase creation probability (*p*_*R*_) and error threshold (*e*_*T*_). Surprisingly, the parameter region of highest replicase fraction per vesicle is distinct from the region with the highest propensity of reaction 9, which occurs for low *e*_*T*_ (see Fig. S12). For high *e*_*T*_, reactions 5-8 collectively dominate over reaction 9 and the non-enzymatic reactions (compare Fig. S12 and S13). Moreover, they use up replicases as catalysts producing parasites, or templates that feed the non-enzymatic reactions (see Fig-1); thereby reducing the propensity of reaction 9 which depends on the presence of both *R* and *T*_*dR*_. For low error-thresholds, the fidelity of replicase creation through reaction 9 is reduced thereby creating a negative feedback loop that reduces the propensity of enzymatic reactions 5-8 and leading to the decrease in number of both RNA and DNA templates. In this regime, the contribution of non-enzymatic reactions to the creation of replicase-encoding DNA template can be significant at high *p*_*R*_ (see Fig. S14 in the electronic supplementary material) and when those reactions are switched off, the average number of *T*_*dR*_ per vesicle drops below 1 and the percentage of vesicles containing *T*_*dR*_ drops below 50% (see Fig. S15 in the electronic supplementary material). Even though reaction 9 is sub-dominant compared to other enzymatic reactions (see Fig. S14(C)), it is essential for ensuring that the average number of replicases inside a vesicle exceeds the number of DNA templates encoding the replicase (see Fig. S16 and section 8 of electronic supplementary material).

## 4 Discussion

Evolution of a protocellular population, aided by the coupling between growth in the number of internal components and division can occur through physical processes only [57, 25]. However, the onset of Darwinian evolution requires heritable and selectively advantageous phenotypes encoded in sequences to be accurately replicated so that they can spread through the population. In this work, we show how the transition to an epoch characterized by Darwinian evolution can be facilitated by the presence of DNA nucleotides. The RNA-templated creation of a DNA sequences using DNA nucleotides lead to new reaction pathways that increase the likelihood of replicase formation through both non-enzymatic and enzymatic channels. Once enzymatic pathways are established, they can help in sustainably creating replicases by increasing both the speed and accuracy of replicase formation. Such enzymatic replicase creation induces symmetry breaking in the number of replicases and replicase-encoding DNA templates, increasing the copy number of the former relative to the latter.

Changing the nature of the protocellular compartment affects the mode of selection and consequently the ability of protocells containing replicases to spread through the population. It is most desirable to get sustained replication and proliferation of a proto-cell containing both the encoded DNA template (*T*_*dR*_) and corresponding replicase (*R*) for low error thresholds and low values of non-enzymatic replicase creation probability. These conditions are satisfied for coacervate and water-in-oil droplet models and we therefore conclude that those compartments will be more effective in ensuring the dominance of protocells containing *R* and *T*_*dR*_ in high and low copy numbers respectively. Although we used coacervates to demonstrate the importance of preferential selection of ribozyme encapsulating protocells on the evolution of the population, those results are applicable for any protocellular compartment that allows for genotype-phenotype coupling. Examples of such coupling have been previously demonstrated for fatty acid vesicles [58, 25].

Our model does not incorporate enzymatic reactions that can create new replicase-encoding DNA templates (*T*_*dR*_) with high fidelity; such templates can be created through non-enzymatic reactions only. That is, most likely, the reason for the observed symmetry-breaking between the number of DNA-encoded replicases (*T*_*dR*_) and replicases (*R*) seen in Fig-4(A) and Fig. S9(A), Fig. S14(A) in the Supplementary Information file. Although we observed sustenance of protocells that contain both *R* and *T*_*dR*_ in the population, such sustenance is possible due to the low degradation rates of *T*_*dR*_ molecules. Hypothetically, the two alternate pathways for creating *T*_*dR*_ enzymatically involves replication of *R* and *T*_*dR*_ with DNA monomers. However, as discussed earlier, it is extremely difficult to replicate complex-folded RNA molecules like the replicase *R*. The alternative pathway involving DNA-templated DNA replication requires the presence of a DNA-dependent DNA-polymerase ribozyme. The 38-6 polymerase ribozyme generated in RNA evolution experiments is far less efficient in catalysing DNA-templated polymerization of DNA sequence using DNA monomers; capable of extending only very short, C-rich primers [33]. Nevertheless, if we assume that the polymerase is also capable of catalysing the creation of new replicase-encoding DNA templates of adequate length, that would necessitate the addition of a new reaction to our model of the form 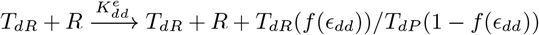.

Simulations including this reaction in the water-in-oil droplet model indicates that the fraction of protocells containing *both R* and *T*_*dR*_ increases significantly compared to the case where the above reaction is absent (*i*.*e*. 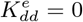). Moreover, in such a situation, the symmetry breaking between the number of replicases and DNA templates encoding replicases also disappears. Whether or not enzymatic synthesis of ribozyme encoding DNA templates would have been possible during the early stages of origin of life, remains an open question for future experiments to address. Hence, a detailed investigation of this new model (with 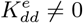) is beyond the scope of this work.

Even though several aspects of our model can be subject to experimental validation, we believe the most compelling test would be to demonstrate the efficacy of auto-catalytic replicase creation through reaction 9 inside a protocell. Although ribozyme-catalysed replication of sequences longer than 100 nucleotides has not yet been experimentally observed, a possible alternative approach might involve ribozyme-catalysed replication of different fragments of a complete DNA template encoding the ribozyme. The RNA fragments generated can then self-assemble to form the whole ribozyme [59, 60, 61], perhaps with the help of auto-catalytic networks of the smaller ribozyme fragments [62]. The versatility of the replicase thus created and its role in catalysing several other reactions, especially in the presence of both RNA and DNA nucleotides, can create a self-sustained chemical system inside a protocell. Although we focused on replicase creation only in this work, other functional ribozymes can also be produced in a similar fashion. We envisage a scenario where multiple functional ribozymes, each encoded by their specific DNA templates, gradually emerge, providing an increasing selective advantage to the protocell encapsulating them and consequently facilitating their proliferation in the population.

The epoch of evolution discussed here predates the Darwinian transition and is also likely to have been of a communal nature [63] characterized by widespread horizontal transfer of sequence elements across protocells. It would be interesting to explore how such horizontal transfer could have driven the evolution towards increasing complexity characterized by the presence of functionally diverse components inside such progenotes. We hope that our work, which establishes a proof of concept for the emergence and proliferation of encoded heritable phenotypes, will motivate the design of novel *in vitro* evolution experiments that will eventually help in further unravelling the mystery of the origin of life.

## 5 Acknowledgments

SR was supported by an INSPIRE graduate fellowship given by SERB, India.

## 6 Authors’ Contributions

S.R.: conceptualization, data curation, formal analysis, investigation, methodology, software, visualization, writing-original draft, writing-review and editing; S.S.: conceptualization, formal analysis, methodology, resources, supervision, writing-original draft, writing-review and editing.

## 7 Data Availability Statement

All codes used to generate the results in the manuscript and in the electronic supplementary material are available at this Github link.

