## Supplementary Information for "The RNA-DNA world and the emergence of DNA-encoded heritable traits"

April 9, 2024

#### Contents

**S1 Experiments on template-directed primer extension** **2**

**S2 Derivation of replication rates** **2**
S2.1 Non-enzymatic template-directed primer extension . . . . . 3
S2.2 Enzymatic template-directed primer extension . . . . . 5

**S3 Dynamical equations** **6**

**S4 Stochastic version of the reaction dynamics** **8**

**S5 Single protocell model: Advantage provided by DNA nucleotides** **9**

**S6 Water-in-Oil droplet model: supplementary figures** **11**

**S7 Coacervate droplet model: supplementary figures** **13**

**S8 Vesicle model** **15**

**S9 Pseudocode for the stochastic model** **18**

#### S1 Experiments on template-directed primer extension

Different experimental groups have demonstrated the efficacy of template-directed primer extension process, both non-enzymatically and in the presence of polymerase ribozymes. Chen et al. (2011) [1] have compared the non-enzymatic primer extension process on both RNA and DNA templates and showed that the mis-incorporation rate during the process is smallest in case of the extension of DNA primer on DNA template. Richert et al. (2013) [2] have shown that the primer extension process has a more complex kinetics. The appearance of a base mismatch during the extension process can increase the downstream mis-incorporation rate of incorrect nucleotides even further. The rates were also shown to depend on the type of activated nucleotides being used for primer extension [3] and the type of cosolutes present in the system [4]. However, these high mutation rates of the non-enzymatic primer extension processes were shown to get reduced significantly when dimer building blocks were used instead of monomers [5, 6, 7], leading to accurate non-enzymatic replication of RNA templates. The enzymatic reaction rates used in this work is derived from the extensive work done by the Joyce group [8, 9, 10] on ribozyme-catalysed primer extension processes. They developed the 24-3 polymerase ribozyme through *in vitro* evolution, which was able to extend primers on a variety of RNA template sequences [8]. The same polymerase was found to act as a reverse transcriptase catalysing the RNA-templated replication of DNA sequences using DNA nucleotides [9]. A mutated version of this ribozyme, the 38-6 polymerase [10] was subsequently shown to be capable of also extending DNA primers on RNA templates and RNA primers on DNA templates. However, extension of DNA primers on DNA templates have not yet been demonstrated.

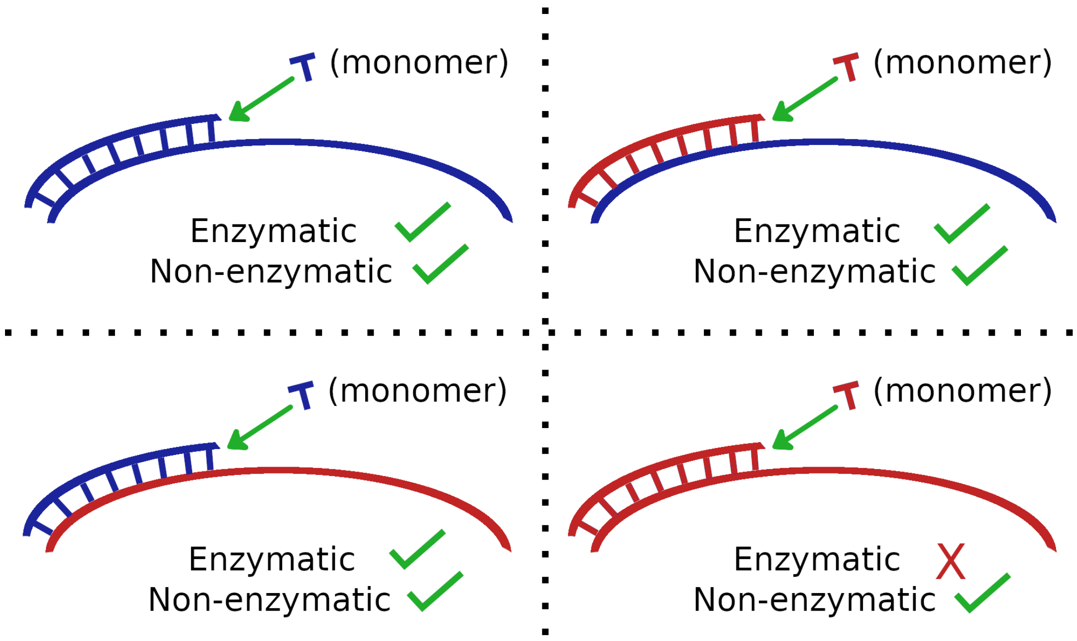


Figure S1: 4 types of template-directed primer extension reactions possible in a mixed RNA-DNA system. Blue denotes RNA strands and RNA monomers. Red denotes DNA strands and DNA monomers. Except for the DNA-templated DNA extension case, all other processes shown can occur both non-enzymatically as well as enzymatically in the presence of replicase ribozymes.

#### S2 Derivation of replication rates

We envisage a scenario where replication occurs via the rolling-circle mechanism [11, 12, 13, 14] using circular templates. The rolling-circle mechanism of replication provides several advantages compared to replication using a linear template. The former mechanism was shown to be more effective in replicating long sequences of the kind considered in our model (200 nucleotides (nt)). It is also capable of evading the strand separation problem that is encountered across a wide range of temperatures in the case of linear templates [15, 16]. Rolling circle replication is a 2-step process. In the first step a small primer attaches to a circular single-stranded RNA or DNA template and extends to full length creating a double stranded circular molecule. In the second step, the fully extended primer extends further by displacing its opposite end from the template. This creates a hanging tail, which gets separated from the template-primer duo when it attains a length similar to the template and becomes a new open-ended strand (see [16] for a pictorial representation of the process). For simplicity, we consider the 2-step rolling circle process as a single step process that leads to the formation of an open-ended RNA/DNA replicate sequence, via template-directed primer extension on a circular template. In order to sustain the rolling circle replication process, circular templates need to be created through some circularization mechanism that converts some of the new open-ended strands into circular strands, either by a ligase ribozyme or by self ligation of the open ends. Here we define good templates as those sequences that fold into simple secondary structures only (for example, hairpin structures). Due to their simple secondary structures, we also assume that their open ends can easily bend and ligate to form circular strands and therefore can sustain the rolling-circle replication process (provided replication can re-create such good templates). For example $A_{50}G_{50}U_{50}C_{50}$ can be a good template. Indeed experiments [17] support the formation of such poly-stack strands in prebiotic environments.

#### S2.1 Non-enzymatic template-directed primer extension

Table S1: Non-enzymatic RNA to RNA Primer extension rates ($h^{-1}$) after a match in the previous step [4]

| Templating nucleotide | Incoming monomer | | | |
|---|---|---|---|---|
| | A | U | G | C |
| A | 0.07 | 0.53 | 0.17 | 0.08 |
| U | 0.92 | 0.12 | 0.38 | 0.017 |
| G | 0.04 | 0.48 | 0.16 | 9.03 |
| C | 0.017 | 0.021 | 1.58 | 0.022 |

Table S2: Non-enzymatic RNA to RNA Primer extension rates ($h^{-1}$) after a mismatch in the previous step [2]

| Templating nucleotide | Incoming monomer | | | |
|---|---|---|---|---|
| | A | U | G | C |
| A | 0.021 | 0.037 | 0.034 | 0.002 |
| U | 0.02 | 0.03 | 0.05 | 0.003 |
| G | 0.018 | 0.03 | 0.04 | 0.005 |
| C | 0.04 | 0.028 | 0.49 | 0.003 |

For deriving the rate of non-enzymatic RNA to RNA replication we used the primer extension rates from [2, 4], where the rate depends on the type of incoming monomer, type of the nucleotide on the template across which the monomer binds and whether there was match or mismatch between the templating base and the nucleotide extending the primer in the preceeding step. Table-S1 and S2 shows the primer extension rates after a match [4] and a mismatch [2] respectively in the previous step, for RNA to RNA replication. Although different experiments [2, 4] have used different activated nucleotides for extending the primers, that can cause a difference in the primer extension rates [3], the complete dataset corresponding to a particular type of activated nucleotides does not exist in the literature. Moreover, our intention here is not to discriminate between different models of primer extension using different types of activated nucleotides. Our main aim is to explore the consequences of template-directed replication of RNA and DNA sequences using both RNA and DNA templates. We therefore ignored the differences due to different types of activated nucleotides and combined the rate-data from different experiments to create the complete dataset.

We took 100 random sequences of length 200 nt and carried out sequence level stochastic simulation of the primer extension process until the primers were fully extended. The average time taken was obtained by averaging over the replication time for the 100 random sequences. The average replication rate was calculated by taking the inverse of the average time for replication.

Table S3: Non-enzymatic DNA to DNA Primer extension rates ($h^{-1}$) after a match in the previous step [2, 18]

| Templating nucleotide | Incoming monomer | | | |
|---|---|---|---|---|
| | A | T | G | C |
| A | 0.125 | 0.843 | 0.125 | 0.125 |
| T | 1.083 | 0.045 | 0.045 | 0.045 |
| G | 0.019 | 0.019 | 0.019 | 1.16 |
| C | 0.035 | 0.035 | 1.11 | 0.035 |

Table S4: Non-enzymatic DNA to DNA Primer extension rates ($h^{-1}$) after a mismatch in the previous step [2, 18]

| Templating nucleotide | Incoming monomer | | | |
|---|---|---|---|---|
| | A | T | G | C |
| A | 0.031 | 0.027 | 0.031 | 0.031 |
| T | 0.036 | 0.028 | 0.028 | 0.028 |
| G | 0.027 | 0.027 | 0.027 | 0.038 |
| C | 0.028 | 0.028 | 0.037 | 0.028 |

Similarly we calculated the rate of non-enzymatic DNA to DNA replication by using the experimentally determined rates [2, 18] (shown in Table-S3 and S4) of non-enzymatic primer extension.

The primer extension rates after a mismatch in the case of non-enzymatic RNA to DNA and DNA to RNA replication were not available and only the rates after a match are known [1] (Table-S5 and Table-S6). Nevertheless mismatches are expected to slow down the rates for these mixed replication processes [2] and the extension rates after a mismatch needs to be appropriately scaled down for these cases (RNA to DNA and DNA to RNA) as well. To do so, we first calculated rates for the RNA to RNA ($K_{rr}^{nm}$) and DNA to DNA ($K_{dd}^{nm}$) replication cases by ignoring the rates after a mismatch ($nm$ in the superscript stands for no mismatch). This allowed us to estimate the factor ($f_{rr}$ and $f_{dd}$) by which reduction of the rates of RNA to RNA ($K_{rr}^{n}$) and DNA to DNA ($K_{dd}^{n}$) replication occurs under the influence of mismatches by calculating the ratio of the complete replication rate obtained by ignoring and incorporating the effect of mismatches respectively. $f_{rr} = K_{rr}^{nm}/K_{rr}^{n}$ and $f_{dd} = K_{dd}^{nm}/K_{dd}^{n}$ are the corresponding reduction factors for RNA to RNA replication and DNA to DNA replication respectively, and we found that their values are nearly equal ($f_{rr} = f_{dd} \approx 2.1$). We then assumed that the slowdown of replication happens by the same factor in the case of RNA to DNA and DNA to RNA replication, from the available rates that do not consider the effect of mismatches. We therefore scaled down the complete replication rate (to replicate 200-mer strands) by that factor to get the reduced replication rate in the presence of mismatches for the latter two cases.

The radius of our model compartment was taken to be $\sim 57\ nm$. 1 template inside such a volume amounts to a template concentration of $2.14\ \mu M$, which is larger than the concentration of $1.3\ \mu M$ used in the experiments. We therefore multiply all 4 of the rate constants with a factor $2.14/1.3$ to get the values presented in Table 1 in the main text.

Table S5: Non-enzymatic RNA to DNA Primer extension rates ($h^{-1}$) after a match in the previous step [1]

| Templating nucleotide | Incoming monomer | | | |
|---|---|---|---|---|
| | A | U | G | C |
| A | 0.016 | 1.6 | 0.52 | 0.022 |
| U | 0.30 | 0.029 | 0.022 | 0.0081 |
| G | 0.034 | 0.75 | 0.25 | 2.5 |
| C | 0.008 | 0.0071 | 1.6 | 0.00085 |

Table S6: Non-enzymatic DNA to RNA Primer extension rates ($h^{-1}$) after a match in the previous step [1]

| Templating nucleotide | Incoming monomer | | | |
|---|---|---|---|---|
| | A | U | G | C |
| A | 0.029 | 0.47 | 0.031 | 0.029 |
| U | 0.093 | 0.064 | 0.065 | 0.019 |
| G | 0.031 | 0.29 | 0.11 | 1.1 |
| C | 0.032 | 0.021 | 3.8 | 0.022 |

##### S2.2 Enzymatic template-directed primer extension

For ribozyme catalyzed replication of RNA strands with RNA nucleotides the replication occurs in two phases. In the first phase replication occurs at a rate $2\ min^{-1}$ up to $20\%$ of the template length (32 nt). In the second phase the replication occurs at a rate $0.073\ min^{-1}$ for the remaining $80\%$ of the template length [9]. We extrapolated this observation to calculate the average replication rate of 200-mer templates in our model, i.e. we got an average replication rate $= 60 \times (40/2 + 160/0.073)^{-1} \sim 0.027\ h^{-1}$ for replication of 200-mer templates. An alternate method involves using the rate $2\ min^{-1}$ up to the length 6 nt on the template, as observed in the experiment for template length of 32 nt and use the slower rate $0.073\ min^{-1}$ for copying the remaining 194 nucleotides of the 200-mer template. This leads to a slightly smaller value of the average replication rate $= 60 \times (6/2 + 194/0.073)^{-1} \sim 0.023\ h^{-1}$, which is still much larger than the non-enzymatic rates. We have verified, by sampling different regions of parameter space, that we get similar results even if we adopt the second procedure for calculating the average replication rate of 200-mer templates.

In the case of enzymatic replication at least 1 template and 1 ribozyme needs to be present inside a compartment, which accounts for a concentration of $2.14\ \mu M$ for both the template and the ribozyme. The template concentration is much larger than the concentration ($125\ nM$) used in the experiment. Hence, according to the Michaelis-Menten equation we can make the replication rate independent of template concentrations if at least one template is present inside a compartment. The ribozyme concentration is also much larger inside our compartment volume. Since the rate is proportional to the ribozyme concentration, we multiply the rate constant with a factor $2.14/0.125$ to get the value presented in the table. Similarly for the case of enzymatic RNA to DNA replication the first phase of replication occurs at a rate $1.1\ min^{-1}$ for up to $35\%$ of the template length. In the second phase the replication occurs at a rate $0.029\ min^{-1}$ for the remaining $65\%$ of the template length [9]. Therefore following the same procedure, we derive the rate constant $= (2.14/0.125) \times 60 \times (70/1.1 + 130/0.029)^{-1} \sim 0.226\ h^{-1}$. Finally, in case of DNA to RNA replication, the longest templating region replicated under the action of the ribozyme was $32\ nt$ long withing $24\ hr$ [10]. This suggests that the replication of a 200-mer strand will occur at a rate of $(200 \times 24/32)^{-1} \sim 0.0067\ h^{-1}$. The ribozyme concentration used in this experiment was $100\ nM$. Therefore we multiply the rate constant with a factor $2.14/0.1$ to get the value presented in Table-1 of the main text.

#### S3 Dynamical equations

The differential equations based on reaction-1 to 10, representing the temporal variation of each type of molecule, are given below. In the following equations $\theta(x) = 1$ for $x >= 1$ and 0 otherwise. The probability modulator function $f(\epsilon_{xy}) = 1/(1 + exp(\epsilon_{xy} - e_T))$. The key variables and parameters are listed in the table below for convenience.

Table S7: Details of each term used in Equ-S1 to S6

| Term | Definition | Value |
|---|---|---|
| $T_r^i$ | Number of RNA templates inside protocell $i$ | variable |
| $P^i$ | Number of RNA parasites inside protocell $i$ | variable |
| $R^i$ | Number of replicases inside protocell $i$ | variable |
| $T_d^i$ | Number of DNA version of templates $T_r$ inside protocell $i$ | variable |
| $T_{dP}^i$ | Number of parasite encoding DNA templates inside protocell $i$ | variable |
| $T_{dR}^i$ | Number of replicase encoding DNA templates inside protocell $i$ | variable |
| $f(\epsilon_{xy})$ | Probability of accurate replication in case of enzymatic replication. xy={rr,rd,dr} | function |
| $\epsilon_{rr}$ | % error in case of ribozyme catalyzed RNA to RNA replication | 8 |
| $\epsilon_{rd}$ | % error in case of ribozyme catalyzed RNA to DNA replication | 3 |
| $\epsilon_{dr}$ | % error in case of ribozyme catalyzed DNA to RNA replication | 3 |
| $e_T$ | Error threshold for ribozyme catalyzed replication | 0% to 8% |
| $p_R$ | Probability of $R/T_{dR}$ creation from non-enzymatic replication | 0.0002 to 0.04 |

$$\dot{T_r^i} = f(\epsilon_{rr})K_{rr}^e\theta(T_r^i)R^i + f(\epsilon_{dr})K_{dr}^e\theta(T_d^i)R^i - h_rT_r^i \quad \text{(S1)}$$

$$\dot{P^i} = (1 - p_R)K_{rr}^nT_r^i + (1 - p_R)K_{dr}^n(T_d^i + T_{dP}^i + T_{dR}^i) + (1 - f(\epsilon_{rr}))K_{rr}^e\theta(T_r^i)R^i + (1 - f(\epsilon_{dr}))K_{dr}^e\theta(T_d^i + T_{dR}^i)R^i + K_{dr}^e\theta(T_{dP}^i)R^i - h_rP^i \quad \text{(S2)}$$

$$\dot{R^i} = p_RK_{rr}^nT_r^i + p_RK_{dr}^n(T_d^i + T_{dP}^i + T_{dR}^i) + f(\epsilon_{dr})K_{dr}^e\theta(T_{dR}^i)R^i - h_rR^i \quad \text{(S3)}$$

$$\dot{T_d^i} = f(\epsilon_{rd})K_{rd}^e\theta(T_r^i)R^i - h_dT_d^i \quad \text{(S4)}$$

$$\dot{T_{dP}^i} = (1 - p_R)K_{rd}^nT_r^i + (1 - p_R)K_{dd}^n(T_d^i + T_{dP}^i + T_{dR}^i) + (1 - f(\epsilon_{rd}))K_{rd}^e\theta(T_r^i)R^i - h_dT_{dP}^i \quad \text{(S5)}$$

$$\dot{T_{dR}^i} = p_RK_{rd}^nT_r^i + p_RK_{dd}^n(T_d^i + T_{dP}^i + T_{dR}^i) - h_dT_{dR}^i \quad \text{(S6)}$$

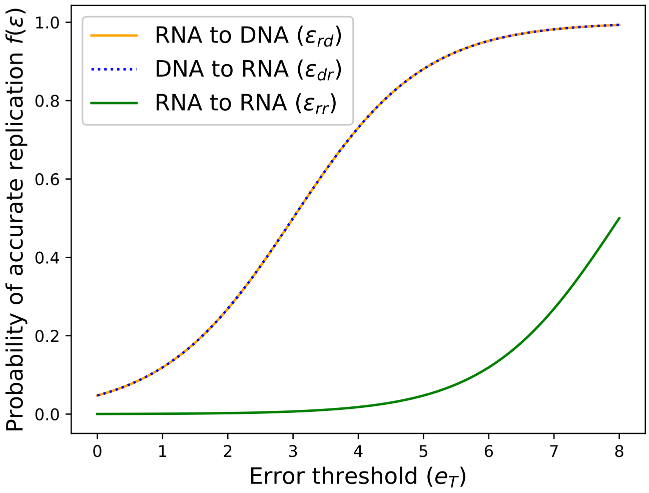


Figure S2: Variation of the probability of accurate enzymatic replication ($f(\epsilon)$) with the error threshold parameter $e_T$. The figure shows probabilities for the cases of RNA to RNA ($f(\epsilon_{rr})$), RNA to DNA ($f(\epsilon_{rd})$) and DNA to RNA ($f(\epsilon_{dr})$) replication.The probability of accurate replication from RNA to DNA ($f(\epsilon_{rd})$) and DNA to RNA ($f(\epsilon_{dr})$) are represented by the solid orange line and dotted blue line respectively. They overlap since $\epsilon_{rd} = \epsilon_{dr}$.

### S4 Stochastic version of the reaction dynamics

In the stochastic version of the reaction dynamics, we calculate total reaction rate for each protocellular compartment and then determine the time step size $dt$ as inverse of the maximum of those rates ($dt = 1/max(tot\_rate)$, where $tot\_rate_i = \sum_{j=1}^{11} Re_i(j)$, with $Re_i(j)$ being the rate of reaction-$j$ inside protocell $i$). At every time step, a protocell can undergo any of the 11 types of reactions with probability $tot\_rate_i \times dt$. The type of reaction that will occur is chosen randomly based on the relative reaction propensities of the 11 types of reactions:

$$Rel_i(1) = K^n_{rr}\, T^i_r/tot\_rate_i,\ Rel_i(2) = K^n_{rd}\, T^i_r/tot\_rate_i$$

$$Rel_i(3) = K^n_{dr}(T^i_d + T^i_{dP} + T^i_{dR})/tot\_rate_i,\ Rel_i(4) = K^n_{dd}(T^i_d + T^i_{dP} + T^i_{dR})/tot\_rate_i$$

$$Rel_i(5) = K^e_{rr}\, R\,\theta(T^i_r)/tot\_rate_i,\ Rel_i(6) = K^e_{rd}\, R\,\theta(T^i_r)/tot\_rate_i$$

$$Rel_i(7) = K^e_{dr}\, R\,\theta(T^i_d)/tot\_rate_i,\ Rel_i(8) = K^e_{dr}\, R\,\theta(T^i_{dP})/tot\_rate_i,\ Rel_i(9) = K^e_{dr}\, R\,\theta(T^i_{dR})/tot\_rate_i$$

$$Rel_i(10) = h_r(T^i_r + P^i + R^i)/tot\_rate_i,\ Rel_i(11) = h_d(T_d + T_{dP} + T_{dR})/tot\_rate_i$$

However, while calculating the relative reaction propensity values for the heatmaps, we divide the individual rates of reaction-1 to 9 with $tot\_rate_i = \sum_{j=1}^{9} Re_i(j)$, i.e. we consider the relative propensity with respect to replication reactions only and disregard the degradation reactions.

For estimating the average fraction of different types of strands per protocell, we carry out both an ensemble average (over $N$ protocells) and time average (over $\tau$ time-steps) of the abundance values per protocell at equilibrium. For example: $Avg(R) = \frac{1}{\tau}\sum_{\nu=t}^{\nu=t+\tau} \frac{1}{N}\sum_{i=1}^{N} R^\nu_i$. Then we calculate the relative values by dividing each value with the sum of average values for all 6 types of strands.

For estimating the average fraction of protocells that contain a particular type of strand at equilibrium, we count the fraction of protocells ($Fr^t(R)$) which contains that type of strand at every time step after equilibrium is reached. Then we take time average of those values: $Fr(R) = \frac{1}{\tau}\sum_{\nu=t}^{\nu=t+\tau} Fr^\nu(R)$.

The time averaging for the water-in-oil-droplet model and coacervate droplet model is done over the interval of 25000 hours to 30000 hours and 40000 hours to 45000 hours respectively, i.e. $\tau = 5000$ hours. However, since the simulations of the vesicle model reaches equilibrium at an earlier time, the time averaging for the vesicle model is done over the interval of 4000 hours to 5000 hours, i.e. $\tau = 1000$ hours.

### S5 Single protocell model: Advantage provided by DNA nucleotides

In order to understand the evolutionary advantage that DNA nucleotides provide, we compared the outcome of reactions in both non-enzymatic and enzymatic reaction networks inside a single protocell in an RNA-DNA world with that of an RNA-only world. In the RNA world, when only non-enzymatic replication is possible, a sub-set of the reaction network (reaction 1) shown in Fig-1 (main text) is relevant. We numerically solved equations S1-S6 (with only terms representing non-enzymatic reactions) inside a single protocell which initially contained 10 RNA templates $T_r$ (see [Fig-S3(A)]) and compared the result with the outcome of allowing for DNA monomers to be present in the system as well (see [Fig-S3(B)]).

The number of strands of all types of molecules gradually decreases after an initial growth spurt, when the replicase ribozyme is not present (Fig-S3(A)). Without enzymatic replication, only reaction-1 can create new strands but growth via reaction-1 is linear and dependent only on the existing RNA templates as it cannot create new templates ($T_r$). Consequently, the initial population of templates is gradually depleted due to the degradation of the RNA templates and the replication process stops completely.

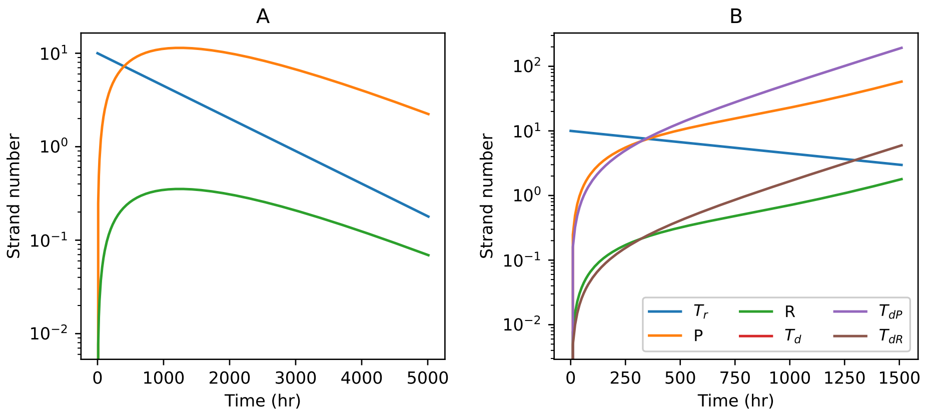


Figure S3: Time evolution of strands when only the non-enzymatic reactions occur inside a single vesicle. The figure shows variation of the number of strands with time **A:** for an RNA only system; **B:** when both RNA and DNA monomers are present.

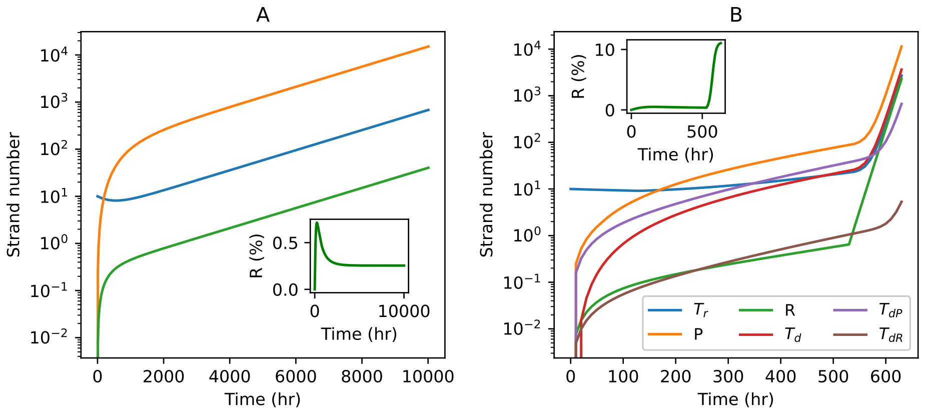


Figure S4: Time evolution of strand numbers inside a single protocell when non-enzymatic as well as enzymatic reactions can occur; for the **A:** RNA only system; **B:** system containing both RNA and DNA monomers. The insets show the time evolution of the percentage of replicase ribozymes among the strand population.

It is clear from Fig-1 (main text) that the presence of DNA nucleotides increases the size of both the non-enzymatic as well as enzymatic reaction networks by opening up new reaction pathways in the non-enzymatic (reactions 2 to 4) and enzymatic (reactions 5 to 9) reaction network. Hence, including DNA nucleotides in the non-enzymatic case leads to an exponential growth in the number of strands (Fig-S3(B)), because Equ. 2 (main text) can create DNA templates $T_{dP}$ and $T_{dR}$ (see Fig-1). Once the DNA templates $T_{dP}$ and $T_{dR}$ are created, they can sustain their own production as well as the production of replicases and parasites in the protocell through reactions 3 and 4, even when the original RNA templates ($T_r$) are degraded.

The chance appearance of a replicase ribozyme, can kick-start enzymatic reactions leading to growth in the number of all possible strands in both the RNA world and the RNA-DNA world (see Fig-S4). From Fig-S4(A) and (B) it is clear that presence of DNA can increase the chances of replicase formation by increasing the number of pathways through which a replicase can be produced. This happens because only reaction-1 can create ribozymes through error-prone replication of RNA templates $T_r$ in an RNA world. However, in presence of DNA monomers, replicase encoding DNA strands ($T_{dR}$) can act as templates for replicase creation both non-enzymatically (through reaction-3) and enzymatically (through reaction-9). Although chance replicase production crucially depends on the probability $p_R$, once present, it ensures faster and more accurate replicase creation which has a positive feedback on other enzymatic reactions, further enhancing the number of replicases inside the protocell (see inset figures in Fig-S4).

#### S6 Water-in-Oil droplet model: supplementary figures

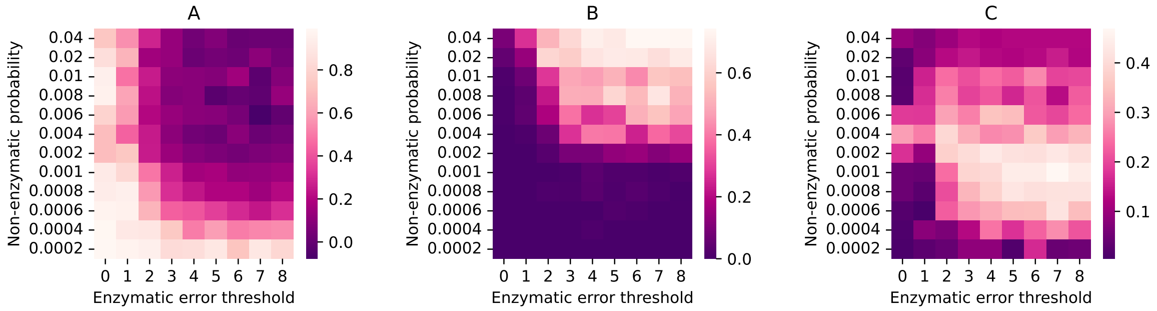


Figure S5: Stochastic simulation of a population of water-in-oil droplets containing strands. Heatmaps showing the variation of **A:** average reaction propensity of the non-enzymatic reactions; **B:** average reaction propensity of reaction-5, 6, 7 together and **C:** average reaction propensity of the reaction-8 per droplet; with different non-enzymatic ribozyme creation probabilities and error thresholds of enzymatic replications. The heatmaps are generated by taking both ensemble average and time average of the quantities at equilibrium.

The non-enzymatic reactions dominate only at lower values of $p_R$ and lower values of error threshold $e_T$ (Fig-S5(A)), because ribozymes are less likely to be created in this parameter region. Even if replicases are created, the enzymatic replication reactions they catalyze will have low fidelity due to low error threshold values. On the contrary, the enzymatic reactions-5, 6 and 7 dominate for higher values of both $p_R$ and $e_T$ (Fig-S5(B)), as more ribozymes can be created at high values of $p_R$ and more templates ($T_r$ and $T_d$) can be re-created by those reactions at high values of $e_T$. The propensity of reaction-8 is highest in the same region where the propensity of the reaction-9 is highest (Fig-S5(C) and Fig-3(C)), i.e. at moderately low values of $p_R$ and moderate to high values of $e_T$. In this region the reaction-9 is becomes the main source of replicase creation and those replicase ribozymes are used to primarily replicate parasitic DNA templates $T_{dP}$ through the reaction-8, leading to their high abundance (see Fig-4(A) of the main text).

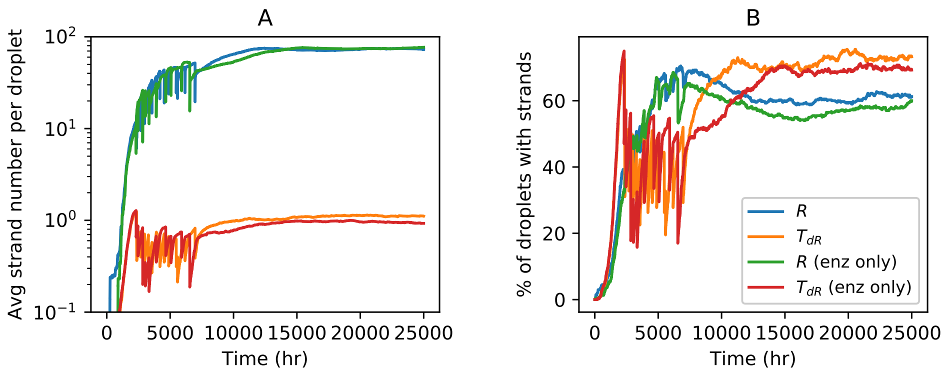


Figure S6: Time evolution when the non-enzymatic reactions are switched off for the water-in-oil droplets inside which both $R$ and $T_{dR}$ emerges ($p_R = 0.002,\ e_T = 4$). Variation of **A:** Average number of $R$ and $T_{dR}$ per droplet (y axis is in logscale); **B:** Percentage of droplets containing $R$ and $T_{dR}$ with time. The standard case that includes the non-enzymatic reactions is also shown for comparison. Panels **A** and **B** have the same legend.

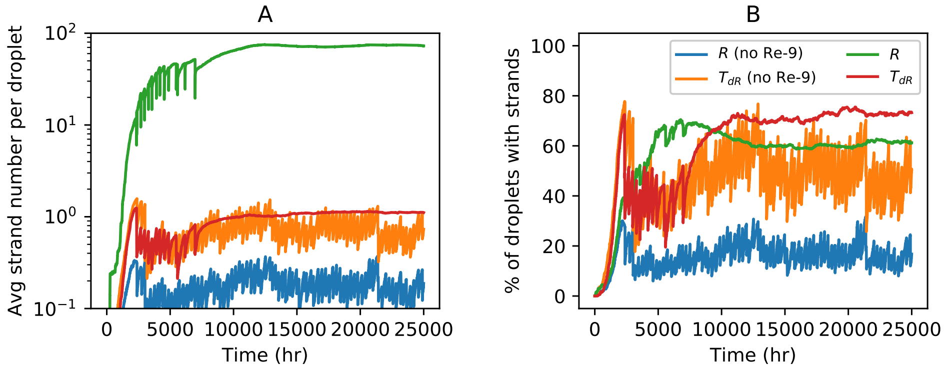


Figure S7: The effect of turning off reaction-9 (enzymatic replicase creation reaction) on the population evolution in the water-in-oil droplet model ($p_R = 0.002,\ e_T = 4$). Time variation of the **A:** average number of $R$ and $T_{dR}$ per droplet (y axis is in logscale); **B:** percentage of droplets containing $R$ and $T_{dR}$. The standard case that includes reaction-9 is also shown for comparison. Panels **A** and **B** have the same legend.

### S7 Coacervate droplet model: supplementary figures

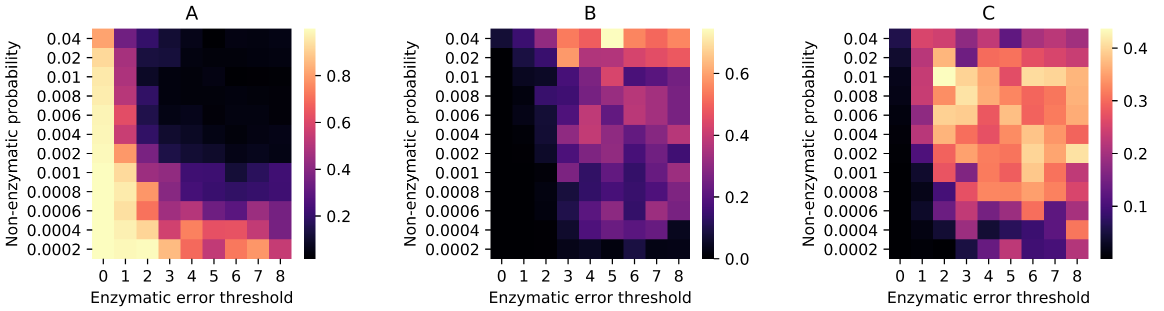


Figure S8: Stochastic simulation of a population of coacervate droplets containing strands. Heatmaps showing the variation of **A:** average reaction propensity of the non-enzymatic reactions; **B:** average reaction propensity of reaction-5, 6, 7 together and **C:** average reaction propensity of the reaction-8 per droplet; with different non-enzymatic ribozyme creation probabilities and error thresholds of enzymatic replications. The heatmaps are generated by taking both ensemble average and time average of the quantities at equilibrium.

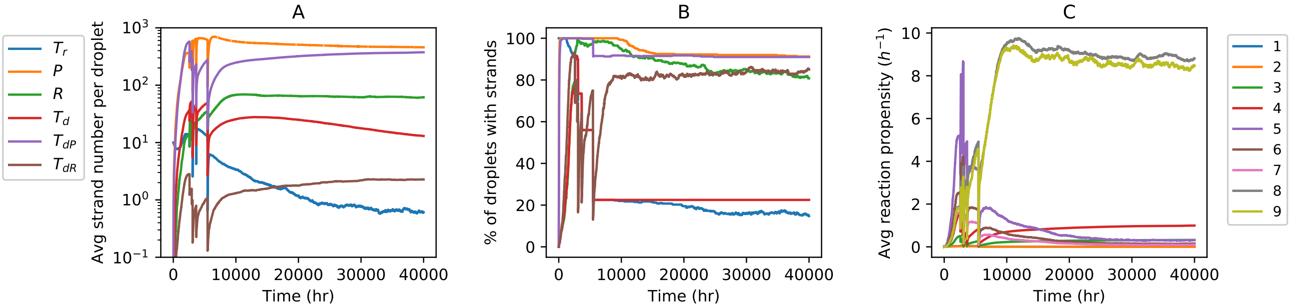


Figure S9: Stochastic simulation of a population of coacervate droplets ($p_R = 0.006,\ e_T = 2$). Time evolution of the **A:** Average number of different types of strands per droplet; **B:** Percentage of droplets containing different types of strands; **C:** Average reaction propensities of the 9 types of reactions per droplet. Panels **A** and **B** have the same legends.

In the case of coacervate droplets, non-enzymatic reactions dominate in the region of low $p_R$ and low $e_T$ values (Fig-S8(A)) for the same reasons as expounded in the water-in-oil droplet scenario. Reactions 5-7 dominate in the upper-right corner where the values of both of these parameters are high (Fig-S8(B)). However, the region of parameter space for their dominance is smaller compared to water-in-oil-droplet model. The propensity of reaction-8 is high in the same region where the propensity of reaction-9 is high (moderate $p_R$) (Fig-S8(C)). As the abundance of $T_{dP}$ is much higher per droplet compared to $T_{dR}$ (Fig-S9(A)), every droplet that contains $T_{dR}$ also contains $T_{dP}$ (Fig-S9(B)). Therefore reactions 8 and 9 occurs with similar propensities when replicases are available, as the propensities of enzymatic reactions depend only on the replicase numbers. The time evolution plots for the percentage of droplets containing different types of molecules (Fig-S9(B)) show that the percentage of droplets containing both $R$ & $T_{dR}$ can be as high as $\sim 80\%$ even for fairly low $p_R$ and low error-thresholds. Even when non-enzymatic reactions are switched off (Fig-S10(A)), the average number of $R$ per droplet remain high ($\mathcal{O}(10)$) with these replicases being present along with $\sim 1\ T_{dR}$ per droplet. Moreover, no significant drop is observed in the number of droplets containing both $R$ & $T_{dR}$ (Fig-S10(B)). Fig-S11 highlights the importance of reaction-9 in driving the formation and proliferation of such droplets. In the absence of reaction-9, which is primarily responsible for efficient replicase creation, the average number of replicases per droplet drops significantly to $\sim 1$ per droplet on par with the average number of $T_{dR}$ per droplet.

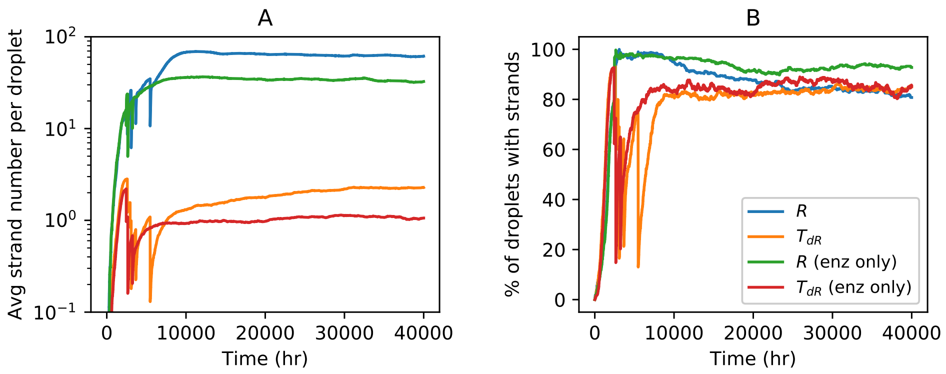


Figure S10: Time evolution when the non-enzymatic reactions are switched off for the coacervate droplets inside which both $R$ and $T_{dR}$ emerges ($p_R = 0.006,\ e_T = 2$). Variation of **A:** Average number of $R$ and $T_{dR}$ per droplet (y axis is in logscale); **B:** Percentage of droplets containing $R$ and $T_{dR}$ with time. The standard case that includes the non-enzymatic reactions is also shown for comparison. Panels **A** and **B** have the same legend.

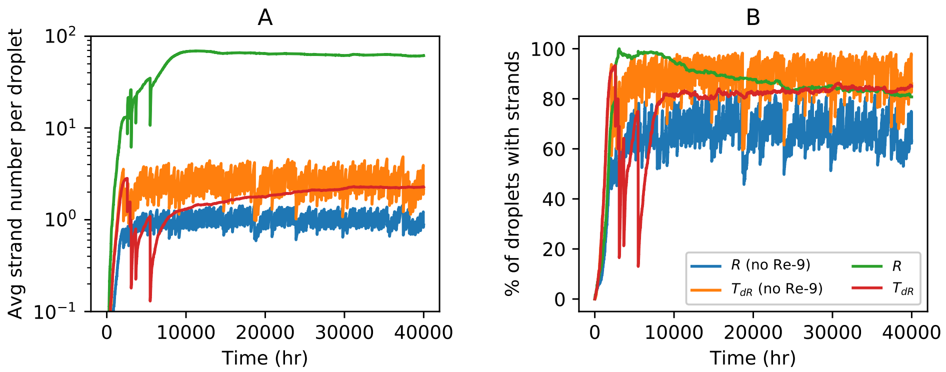


Figure S11: The effect of turning off reaction-9 (enzymatic replicase creation reaction) on the population evolution in the coacervate droplet model ($p_R = 0.006,\ e_T = 2$). Time variation of the **A:** average number of $R$ and $T_{dR}$ per droplet (y axis is in logscale); **B:** percentage of droplets containing $R$ and $T_{dR}$. The standard case that includes reaction-9 is also shown for comparison. Panels **A** and **B** have the same legend.

### S8 Vesicle model

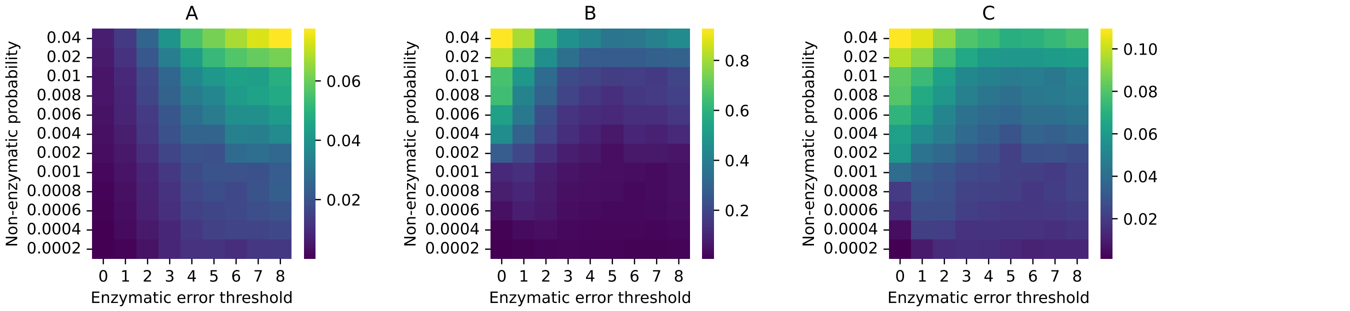


Figure S12: Stochastic simulation of a population of vesicles containing strands. Heatmaps showing the variation of **A:** average fraction of replicase ribozymes per vesicle; **B:** fraction of vesicles containing both replicase ($R$) and replicase encoding DNA template ($T_{dR}$); **C:** average reaction propensity of reaction 9 per vesicle; with different non-enzymatic ribozyme creation probabilities and error thresholds of enzymatic replications. The heatmaps are generated by taking both ensemble average and time average of the quantities at equilibrium.

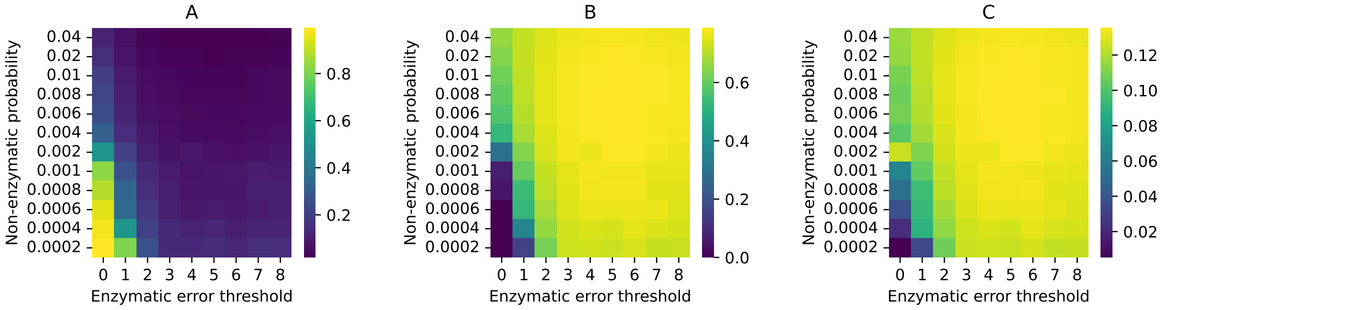


Figure S13: Stochastic simulation of a population of vesicles containing strands. Heatmaps showing the variation of **A:** average reaction propensity of the non-enzymatic reactions; **B:** average reaction propensity of reaction-5, 6, 7 together and **C:** average reaction propensity of the reaction-8 per vesicle; with different non-enzymatic ribozyme creation probabilities and error thresholds of enzymatic replications. The heatmaps are generated by taking both ensemble average and time average of the quantities at equilibrium.

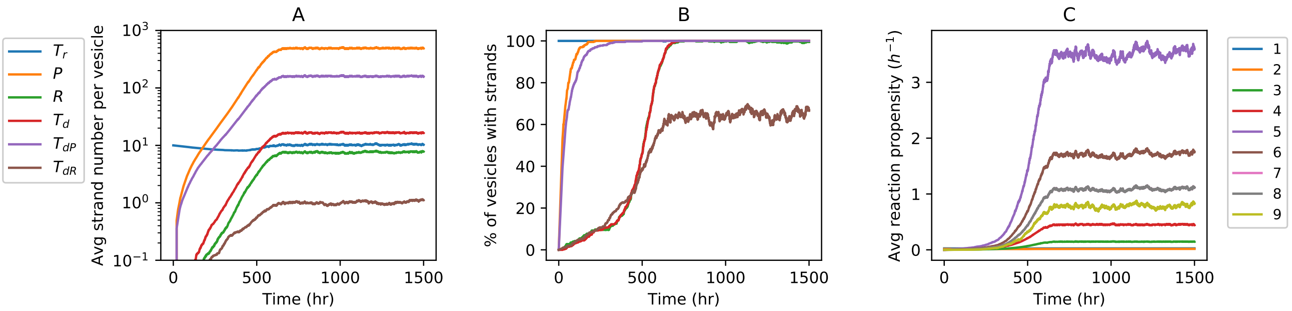


Figure S14: Stochastic simulation of a population of vesicles ($p_R = 0.02,\ e_T = 1$). Time evolution of the **A:** Average number of different types of strands per vesicle; **B:** Percentage of vesicle containing different types of strands; **C:** Average reaction propensities of the 9 types of polymerization reactions per vesicle. Panels **A** and **B** have the same legends.

The near overlap between the trajectory of reaction-7 and 8 in Fig-S14 can explained as follows. Initially the number of vesicles with $T_{dP}$ increases rapidly since the non-enzymatic DNA replication reactions using RNA templates are more likely to create DNA parasitic strands (Fig-S14(B)). The number of vesicles with $R$ grow at a relatively slower rate. However, the emergence of $R$ in the vesicles soon induces the replicase catalysed creation of $T_d$ (through reaction-6). Reaction-7 starts occurring in Vesicles containing both $T_d$ and $R$. Since almost all of the vesicles contain $T_{dP}$ by this time, reaction-8 also starts taking place in these vesicles as well. Even though reaction-7 initially lags behind reaction-8 slightly since the former depends on enzymatic creation of $T_d$, once $T_d$ is created inside all of the vesicles (Fig-S14(B)), the propensities of reactions 7 & 8 become equal as they depend on the number of $R$ only provided there is at least one DNA template of both types. Therefore the time variation curves for reaction-7 and 8 are nearly identical for the vesicle model.

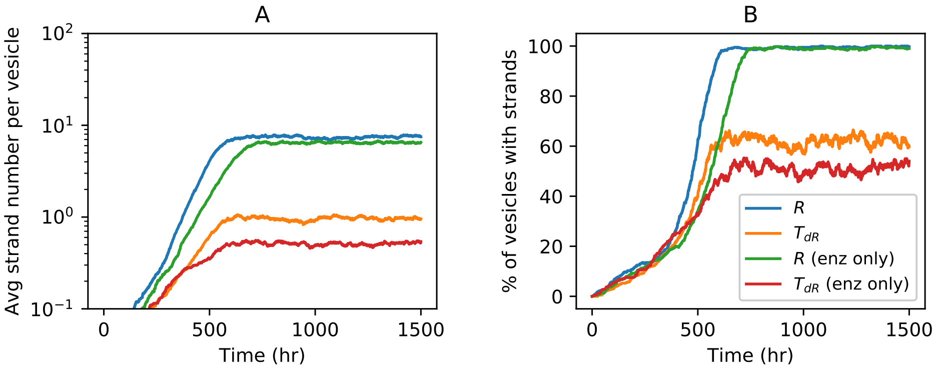


Figure S15: Time evolution when the non-enzymatic reactions are switched off for the vesicles inside which both $R$ and $T_{dR}$ emerges ($p_R = 0.02,\ e_T = 1$). Variation of **A:** Average number of $R$ and $T_{dR}$ per vesicle (y axis is in logscale); **B:** Percentage of vesicles containing $R$ and $T_{dR}$ with time. The standard case that includes the non-enzymatic reactions is also shown for comparison. Panels **A** and **B** have the same legend.

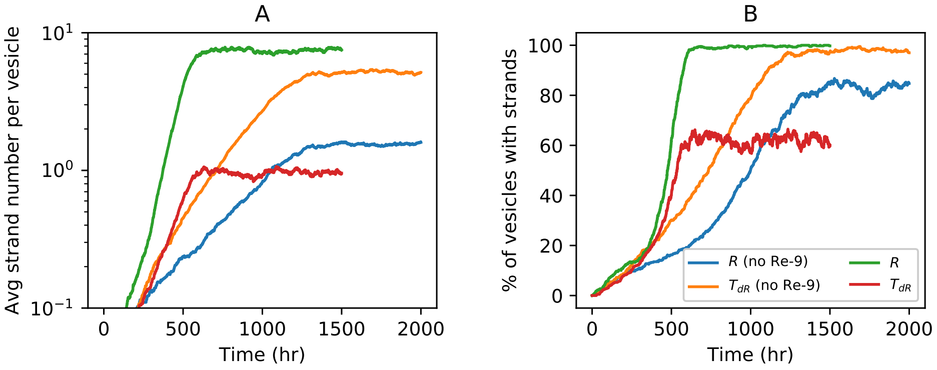


Figure S16: The effect of turning off the reaction-9 (enzymatic replicase creation reaction) on the population evolution in the vesicle model ($p_R = 0.02,\ e_T = 1$). Time variation of the **A:** average number of $R$ and $T_{dR}$ per vesicle (y axis is in logscale); **B:** percentage of vesicles containing $R$ and $T_{dR}$. The standard case that includes reaction-9 is also shown for comparison. Panels **A** and **B** have the same legend.

The reverse asymmetry between the number of $R$ and $T_{dR}$ (i.e. $T_{dR} > R$) in the absence of reaction-9 for a vesicle system occurs because turning off this reaction eliminates a major source of replicase creation. When there is a shortage of replicase ribozymes, the propensities of all enzymatic reactions become smaller and the non-enzymatic reactions promote $T_{dR}$ creation via multiple reaction pathways (reactions 2 and 4 in Fig-1 of the main text). In contrast, non-enzymatic creation of $R$ is possible only through reaction 3. Since, in this case, selection prefers a vesicular compartment having more number of strands, without discriminating between the nature of strands, the vesicles tend to contain more $T_{dR}$ than $R$. Moreover, since $T_{dR}$ has a higher lifetime than $R$, due to its smaller degradation rate, it survives longer inside a vesicle, all of which lead to vesicles where the number of $T_{dR}$ is larger than $R$. Similar results were also obtained in the single protocell model (Fig-S3(B)), in the absence of enzymatic reactions.

#### S9 Pseudocode for the stochastic model

**Initial setup:**

Number of compartments $N = 400$, upper limit for the total number of strands per compartment $V = 1000$
$K^n_{rr} = K^n_{dd} = 0.00255,\ K^n_{rd} = 0.001645,\ K^n_{dr} = 0.000823$
$K^e_{rr} = 0.46447,\ K^e_{rd} = 0.226,\ K^e_{dr} = 0.1434$
$h_r = 0.0008,\ h_d = 0.00008$
Define a function $\theta(x) = 1,\ for\ x > 0$ and $\theta(x) = 0,\ for\ x = 0$
Create a list named cells, under which there are N empty lists denoting each compartment
Fill each cell with a randomly drawn value from the range $(0.5\,p_R, 1.5\,p_R)$,
followed by 6 empty lists, which will contain the 6 types of strands in our model
Create a list of length N, named tot_rate for storing the total rates of each cell

**Time evolution:**

Begin loop over time

- **Determine time step size:**
- Begin loop over N cells
  - Find out the number of different types of strands in the present cell
  - Create a list containing the rates for 11 types of reactions:

$$a = \Big[0.00255\,T^i_r,\ 0.001645\,T^i_r,\ 0.000823(T^i_d + T^i_{dP} + T^i_{dR}),\ 0.00255(T^i_d + T^i_{dP} + T^i_{dR}),$$
$$0.46447\,R\,\theta(T^i_r),\ 0.226\,R\,\theta(T^i_r),\ 0.1434\,R\,\theta(T^i_d),\ 0.1434\,R\,\theta(T^i_{dP}),\ 0.1434\,R\,\theta(T^i_{dR})$$
$$0.0008(T^i_r + P^i + R^i),\ 0.00008(T^i_d + T^i_{dP} + T^i_{dR})\Big]$$

  - If the total number of strands in this cell ≥ V, then make the rates of all 9 types of replication reactions 0
  - Assign tot_rate(i)=sum(a)
- End loop over cells
- Calculate $dt = 1/max(tot\_rate)$
- Change time $t = t + dt$

  **Reactions:**

- Begin loop over cells
  - If $tot\_rate(i) > 0$ and $random.uniform(0, 1) < tot\_rate(i)/max(tot\_rate)$
    * Create a list containing the rates for 11 types of reactions:

$$a = \Big[0.00255\,T^i_r,\ 0.001645\,T^i_r,\ 0.000823(T^i_d + T^i_{dP} + T^i_{dR}),\ 0.00255(T^i_d + T^i_{dP} + T^i_{dR}),$$
$$0.46447\,R\,\theta(T^i_r),\ 0.226\,R\,\theta(T^i_r),\ 0.1434\,R\,\theta(T^i_d),\ 0.1434\,R\,\theta(T^i_{dP}),\ 0.1434\,R\,\theta(T^i_{dR})$$
$$0.0008(T^i_r + P^i + R^i),\ 0.00008(T^i_d + T^i_{dP} + T^i_{dR})\Big]$$

    * If the total number of strands in this cell ≥ V, then make the rates of all 9 types of replication reactions 0

* Choose the type of reaction randomly based the relative propensities ($a/tot\_rate(i)$) of the reactions
* If chosen reaction is 1
  · If $random.uniform(0, 1) < p_R$ add 1 strand to the list of $R$
  · Else add 1 strand to the list of $P$
* If chosen reaction is 2
  · If $random.uniform(0, 1) < p_R$ add 1 strand to the list of $T_{dR}$
  · Else add 1 strand to the list of $T_{dP}$
* If chosen reaction is 3
  · If $random.uniform(0, 1) < p_R$ add 1 strand to the list of $R$
  · Else add 1 strand to the list of $P$
* If chosen reaction is 4
  · If $random.uniform(0, 1) < p_R$ add 1 strand to the list of $T_{dR}$
  · Else add 1 strand to the list of $T_{dP}$
* If chosen reaction is 5
  · If $random.uniform(0, 1) < f(8)$ add 1 strand to the list of $T_r$
  · Else add 1 strand to the list of $P$
* If chosen reaction is 6
  · If $random.uniform(0, 1) < f(3)$ add 1 strand to the list of $T_d$
  · Else add 1 strand to the list of $T_{dP}$
* If chosen reaction is 7
  · If $random.uniform(0, 1) < f(3)$ add 1 strand to the list of $T_r$
  · Else add 1 strand to the list of $P$
* If chosen reaction is 8, add 1 strand to the list of $P$
* If chosen reaction is 9
  · If $random.uniform(0, 1) < f(3)$ add 1 strand to the list of $R$
  · Else add 1 strand to the list of $P$
* If chosen reaction is 10, randomly select a strand from the combined list of $T_r$, $P$ & $R$, then remove that strand
* If chosen reaction is 11, randomly select a strand from the combined list of $T_d$, $T_{dP}$ & $T_{dR}$, then remove that strand

- End loop over cells

**Cell division: vesicle**

- Create a list of cell indices
- Start loop over cells
  – Count total number of strands ($T_r^i + P^i + R^i + T_d^i + T_{dP}^i + T_{dR}^i$) inside the cell
  – If $(T_r^i + P^i + R^i + T_d^i + T_{dP}^i + T_{dR}^i) \geq V$
    * Create an empty list for weakness
    * Start loop over cells
      · If cell index is not marked 'X' and total number of strands inside it ($T_r^j + P^j + R^j + T_d^j + T_{dP}^j + T_{dR}^j$) is lower than that of the dividing cell, then Assign weakness $w_j = (T_r^i + P^i + R^i + T_d^i + T_{dP}^i + T_{dR}^i) - (T_r^j + P^j + R^j + T_d^j + T_{dP}^j + T_{dR}^j)$
      · Else assign weakness $w_j = 0$
    * Choose a cell randomly with probabilities equal to the relative weaknesses of the cells

- * Make the chosen cell empty
  * Mark the cell index for both the dividing cell and the eliminated cell 'X'
  * Start loop over each strand of the dividing cell
    · Draw a random number p
    · If $p < 0.5$ move the strand to the empty cell and remove from the present cell
    · Else keep the strand in the present cell
  * End loop over strands

- End loop over cells

**Cell division: water-in-oil and coacervate droplet**

- If total number of strands is $\geq V/5$ for all cells
  - Randomly shuffle the cell locations in the list cells
  - Start loop over cells
    * In case of coacervate droplets calculate number of $R$ and other types of molecules in cell $i$ and $i+1$
    * Calculate fitness as $f_i = \frac{R^i}{T_r^i+P^i+T_d^i+T_{dP}^i+T_{dR}^i}$ and $f_{i+1} = \frac{R^{i+1}}{T_r^{i+1}+P^{i+1}+T_d^{i+1}+T_{dP}^{i+1}+T_{dR}^{i+1}}$
    * If both $f_i$, $f_{i+1} = 0$ then assign fitness as $f_i = (T_r^i + P^i + R^i + T_d^i + T_{dP}^i + T_{dR}^i)$ and $f_{i+1} = (T_r^{i+1} + P^{i+1} + R^{i+1} + T_d^{i+1} + T_{dP}^{i+1} + T_{dR}^{i+1})$
    * If $random.uniform(0,1) < f_{i+1}/f_i$ then make cell $i$ empty, else make cell $i+1$ empty
    * In case of water-in-oil droplets just make cell $i+1$ empty
    * Start loop over each strand of the dividing cell
      · Draw a random number p
      · If $p < 0.5$ move the strand to the empty cell and remove from the present cell
      · Else keep the strand in the present cell
    * End loop over strands
  - End loop over cells

End loop over time
